# Ghost admixture in eastern gorillas

**DOI:** 10.1101/2022.12.19.521012

**Authors:** Harvinder Pawar, Aigerim Rymbekova, Sebastian Cuadros, Xin Huang, Marc de Manuel, Tom van der Valk, Irene Lobon, Marina Alvarez-Estape, Marc Haber, Olga Dolgova, Sojung Han, Paula Esteller-Cucala, David Juan, Qasim Ayub, Ruben Bautista, Joanna L. Kelley, Omar E. Cornejo, Oscar Lao, Aida M. Andrés, Katerina Guschanski, Benard Ssebide, Mike Cranfield, Chris Tyler-Smith, Yali Xue, Javier Prado-Martinez, Tomas Marques-Bonet, Martin Kuhlwilm

**Author notes:** Correspondence to (M.K.), (T. M.-B.). These authors contributed equally (Tomas Marques-Bonet, Martin Kuhlwilm).

## Abstract

Archaic admixture has had a significant impact on human evolution with multiple events across different clades, including from extinct hominins such as Neanderthals and Denisovans into modern humans. Within the great apes archaic admixture has been identified in chimpanzees and bonobos, but the possibility of such events has not been explored in other species. Here, we address this question using high-coverage whole genome sequences from all four extant gorilla subspecies, including six newly sequenced eastern gorillas from previously unsampled geographic regions. Using Approximate Bayesian Computation (ABC) with neural networks to model the demographic history of gorillas, we find a signature of admixture from an archaic ‘ghost’ lineage into the common ancestor of eastern gorillas, but not western gorillas. We infer that up to 3% of the genome of these individuals is introgressed from an archaic lineage that diverged more than 3 million years ago from the common ancestor of all extant gorillas. This introgression event took place before the split of mountain and eastern lowland gorillas, likely more than 40 thousand years ago, and may have influenced perception of bitter taste in eastern gorillas. When comparing the introgression landscapes of gorillas, humans and bonobos, we find a consistent depletion of introgressed fragments on the X chromosome across these species. However, depletion in protein-coding content is not detectable in eastern gorillas, possibly as a consequence of stronger genetic drift in this species.

## Introduction

Gorillas are a member of the great apes, and form a sister clade to *Homo* (human) and *Pan* (chimpanzees and bonobos). Extant gorillas consist of four recognised subspecies, which cluster into two species, a western species of western lowland (*Gorilla gorilla gorilla*) and Cross River (*Gorilla gorilla diehli)* gorillas and an eastern species of eastern lowland (*Gorilla beringei graueri*) and mountain gorillas (*Gorilla beringei beringei)*^1^. All gorilla subspecies are either endangered or critically endangered under IUCN criteria^2–4^.

The subspecies are distributed across western and eastern Africa in a non-continuous manner (Fig. 1A). The current geographic ranges of the different subspecies differ by size, continuity and ecology, impacting connectivity and population sizes^5^. Western lowland gorillas are endemic to a largely continuous range of considerable size, whereas the other subspecies have much more fragmented distributions^6^. Likewise, western lowland gorillas exhibit the highest genetic diversity of the subspecies^5,7,8^, indicative of long-term high effective population sizes, while eastern gorilla effective population sizes are smaller^9^. Mountain gorillas are currently isolated in two discrete areas, the Virunga Volcanoes Massif and the Bwindi Impenetrable National Park. The Bwindi National Park is located at a lower elevation than the Virunga Volcanoes, and as such exhibits warmer temperatures^10,11^. Previous studies of the demographic history of gorillas did not incorporate information from all subspecies and were not conclusive, especially regarding the divergence time between the eastern and western clade^9,12–15^. This might be due to gene flow from unsampled lineages, which is likely widespread but is often not sufficiently considered in evolutionary studies^16,17^. While uncovering such hidden introgression events in gorillas is not possible from ancient DNA from fossil remains, as has been performed in humans^18^, it is possible to address such questions using genomic data from present-day individuals^19-22^.

**Fig 1:**
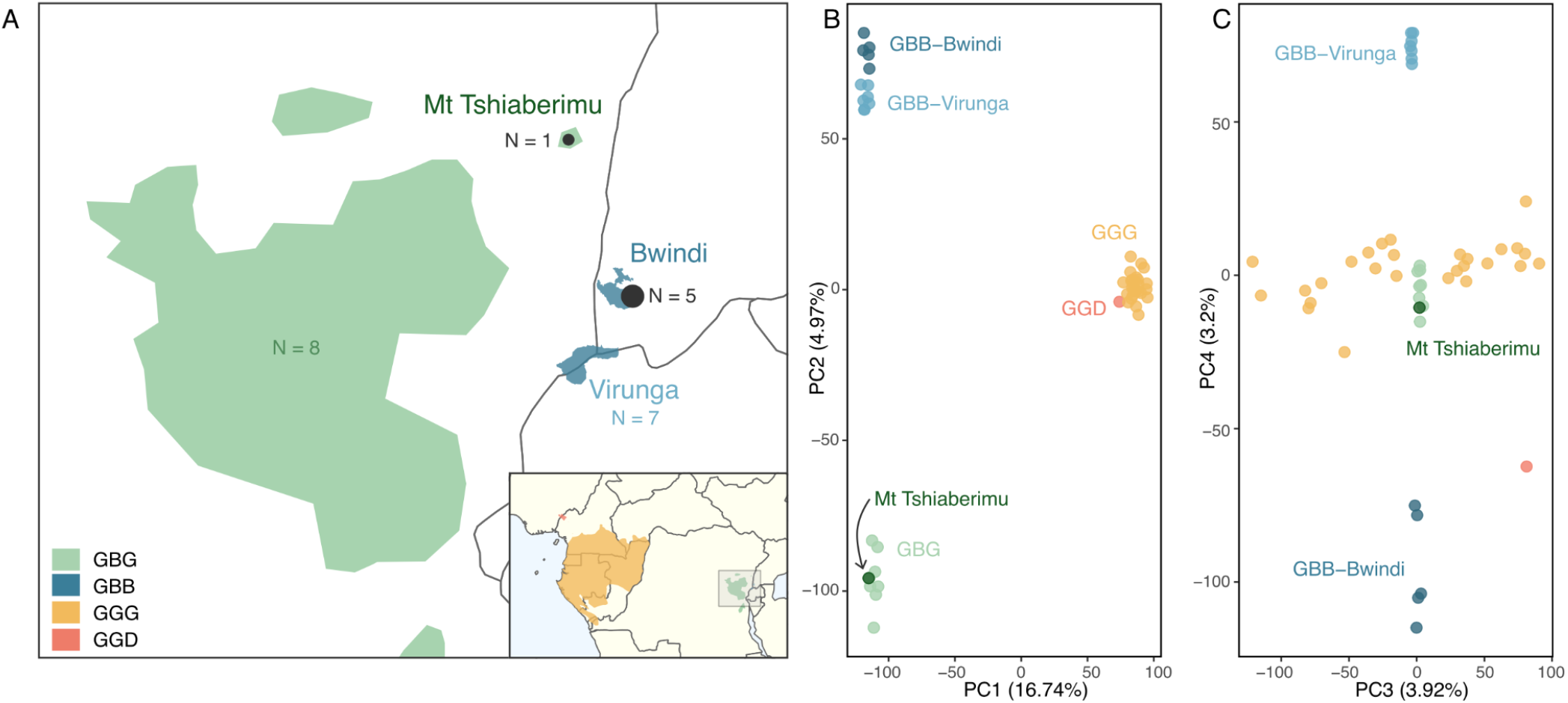
Gorilla samples used in this study. **A** Present geographic distribution of eastern gorillas, with that of the four gorilla subspecies shown in the inset. The newly sequenced samples are given in black, numbers of previously sequenced eastern gorillas are given in colour. GBG = *Gorilla beringei graueri* (Eastern lowland gorilla, n=9); GBB = *Gorilla beringei beringei* (Mountain gorilla, n=12); GGG = *Gorilla gorilla gorilla* (Western lowland gorilla, n=27); GGD = *Gorilla gorilla diehli* (Cross River gorilla, n=1). **B** Principal Component Analysis (PCA) with principal components 1 and 2 shown. **C** PCA with principal components 3 and 4 shown.

To address this question, we use high-coverage whole genome sequences of 28 western and 21 eastern gorillas. In addition to previously published genomes^7,8^, we sequenced the genomes of five mountain gorillas from the Bwindi National Park, and one eastern lowland gorilla from the isolated population of Mount Tshiaberimu. These new genomes contribute to a more complete representation of the genomic diversity of eastern gorillas. Using this expanded dataset, representing all four known gorilla subspecies, we explored the demographic history of gorillas and specifically the hypothesis of ghost introgression, defined as gene flow from an unsampled archaic lineage. Given its significant impact in their sister taxa of *Pan* and *Homo* as well as many other species^18,21,22^, such ghost introgression events may explain some of the uncertainties in previous demographic models for gorillas. Using an Approximate Bayesian Computation (ABC) approach, we find evidence for introgression from an extinct lineage into the common ancestor of eastern gorillas, and characterise some of the functional consequences of this introgressed genetic material.

## Results

### Eastern gorillas form two population clusters

We newly sequenced six eastern gorillas to high coverage (on average, 28.6X). After reprocessing the sequencing data from previous studies (Methods), we obtained a dataset of 49 individuals, with 27 western lowland gorillas, one Cross River gorilla, 12 mountain gorillas and nine eastern lowland gorillas (Table S1). We performed a principal component analysis (PCA, Methods) to ascertain whether the newly sequenced individuals cluster with individuals from the same subspecies. The first PC separates western and eastern gorillas, as previously observed, and the second PC separates mountain gorillas from eastern lowland gorillas (Fig 1B). Since the new individual from the isolated Mount Tshiaberimu population clusters within the distribution of the other eastern lowland gorillas (Fig 1B), this individual is, as expected, considered a representative of this subspecies. The third PC reflects population stratification within western lowland gorillas, whereas the fourth PC separates the eastern gorillas, with the two mountain gorilla populations from Virunga and Bwindi at the extremes (Fig 1C), explaining 3.2% of the variance. This is well in agreement with previous studies^7,8^.

## Demographic modelling favours a ghost lineage in eastern gorillas

To infer a demographic model for the four extant gorilla subspecies, we used a neural-network based Approximate Bayesian Computation (ABC) modelling strategy using windowed summary statistics and extensive simulations (Methods), based on a previous implementation in the *Pan* clade^22^. A main improvement is the implementation of a broad range of informative summary statistics (Table S6), as is common practice for ABC studies in modern and ancient humans^23^.

We first established a demographic null model of the four populations (Fig. S6A,S7), based on previous studies^3,7,8,24–27^. Notably, although none of these studies incorporated whole-genome data from all subspecies, our inferred parameters are largely coherent with previous work (Table S2-S3). Nevertheless, unaccounted demographic events such as ancient population structure or ghost admixture could affect parameter estimates^16^, particularly given evidence in other great apes^22^. Initial exploratory analyses with f4-statistics and admixture graphs (SI 2) did not show any asymmetries between the four gorilla terminal populations, which would arise if ghost admixture had occurred in any of the individual subspecies. However, this does not exclude the possibility of ghost admixture into the common ancestor of eastern or western gorillas, which these methods cannot assess. To account for this and explicitly test if ghost admixture could improve the inferred null demographic model (model A), we considered two more complex demographic models, in which we added the possibility of ‘ghost’ introgression into the common ancestor of eastern gorillas (model B) and western gorillas (model C). We assessed the robustness of our ghost models B and C using a wider parameter space (Fig S11-S13, see Methods), resulting in coherent posteriors with those observed in models B and C (Table S3), albeit with wider confidence intervals, as expected given the increased model complexity (Fig S11). We performed a formal comparison of these models (see Methods), to determine which fits the empirical data best. Model B, with archaic gene flow to the common eastern ancestor had the highest posterior model probability of 0.9973, compared to models A (0.0027) and C (0), and a substantially higher Bayes factor (374, vs. 0.0027 for model A and 0 for model C). In a cross-validation analysis, the model with archaic introgression into eastern gorillas was clearly distinguishable from the model without (Table S4). We conclude that a model with archaic introgression into the common eastern ancestor best explains the observed summary statistics in the empirical data.

We infer that eastern gorillas experienced bottlenecks and generally had lower effective population sizes than western gorillas, while mountain gorillas and eastern lowland gorillas experienced a particularly strong population decrease (Tables S2-S3), as described previously^8,24^. We infer that the eastern subspecies split at 15 thousand years ago (kya) (14-16 kya, 95% credible interval (CI), Tables S2-S3). In agreement with previous studies^13^, we see a population expansion in western lowland gorillas ∼40 kya. Our null demographic model infers a large ancestral population size for western gorillas (Ne=98,135), in comparison to that of other gorilla populations, as well as a split between the western gorilla subspecies at ∼454 kya (448-456 kya CI). Considering that not all summary statistics could be calculated for Cross River gorillas (where only one sample was available), and gene flow between western gorilla subspecies was not modelled, we caution that the confidence in this split time might be low. Finally, we infer that gorillas diverged into two species ∼965 kya (729-1,104 kya CI), which is within the higher range of previous estimates^9,12,28^.

For simplicity, we modelled extant admixture as single migration pulses over one generation, finding a small contribution of gene flow from the common eastern ancestor to the western lowland gorillas of 0.80% (0.06% - 2.14% CI), as well as from western lowland gorillas to the common eastern ancestor of 0.27% (0.22% - 0.43% CI). We infer a contribution of 2.47% of gene flow from an archaic source into the common ancestor of eastern gorillas, with a narrow 95% CI of 2.38-2.49% (Fig 2C). We infer this ghost population diverged from the extant gorilla lineages ∼3.4 million years ago (Mya) (2.98-3.8 Mya, 95% CI). We estimate the timing of this ghost gene flow to have occurred 38,281 years ago, although the CIs for this parameter are wide (22-108 kya, 95% CI) (Fig 2A,2C). By contrast, the posterior distributions for the archaic introgression proportion and the gorilla-ghost divergence time have narrow credible intervals, indicating a strong support with clear peaks for these parameters (Fig 2C). In contrast, our ABC analysis of model C does not confidently infer a contribution of a deeply divergent external lineage into the common ancestor of western gorillas. Instead, the best fit of this model suggests a 0.17% (0.09%-0.4%, 95% CI) contribution from an external lineage at ∼1.1 Mya into the common ancestor of all extant gorillas (Table S3). This marginal contribution is inferred to originate from an external lineage which diverged from extant gorillas 1.9 Mya (1.5 - 3 Mya).

**Fig 2:**
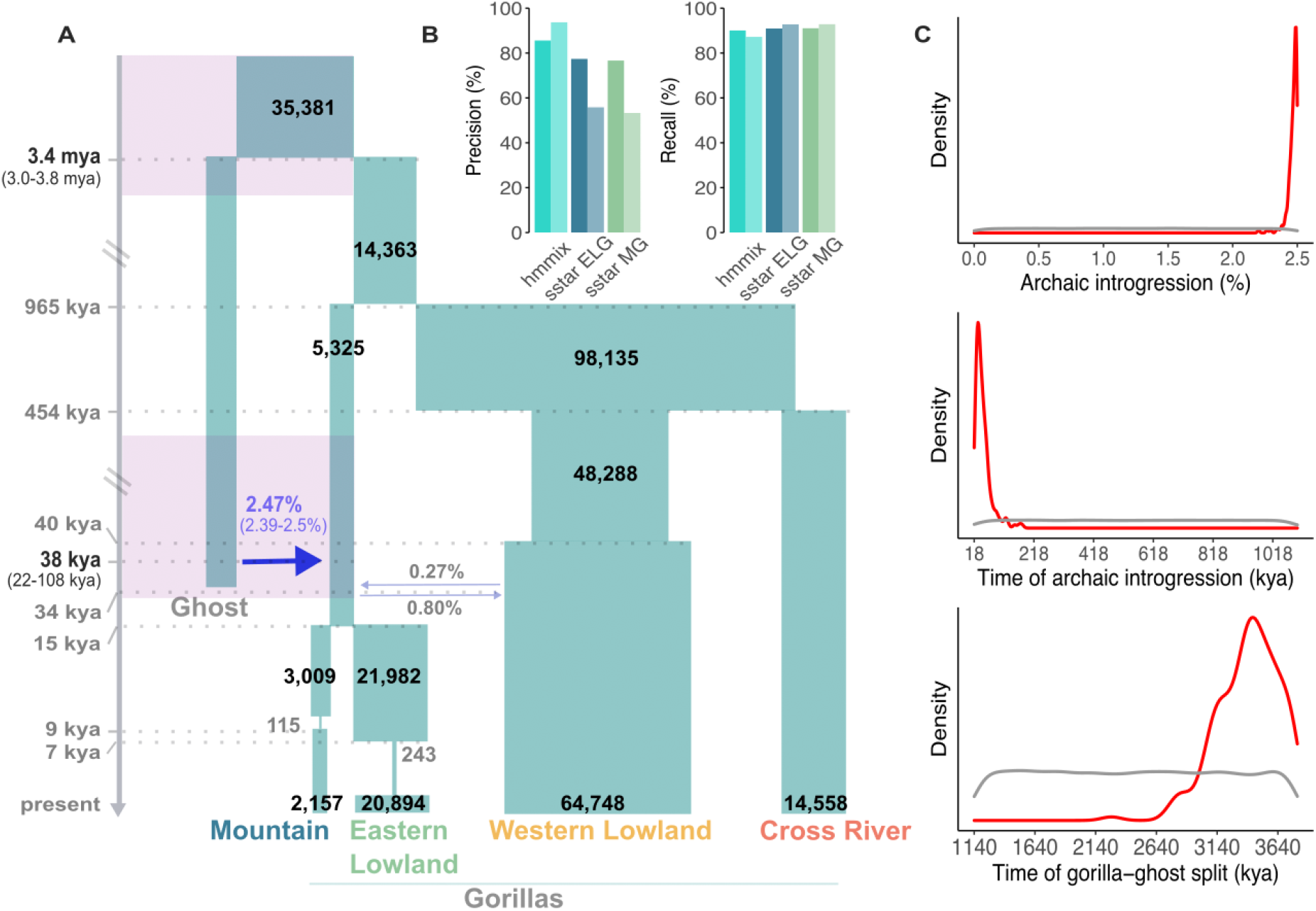
ABC-based demographic model. **A** Model of gorilla population history with archaic admixture from an unsampled ‘ghost’ lineage into the common ancestor of eastern gorillas. 95% credible intervals are shown for the archaic introgression proportion, timing of archaic introgression and archaic divergence (purple timeframes), inferred under ABC modelling. Numbers on blocks represent effective population sizes. kya = thousand years ago; mya = million years ago. **B** Precision and recall of hmmix (at the 95% posterior probability cutoff) and *S*\* (at the 99% quantile using sstar^29^) in simulated data using msprime. Precision (percentage of recovered introgressed fragments) and recall (percentage of true among inferred introgressed fragments) for hmmix and *S\** (for ELG = eastern lowland gorilla and MG = mountain gorilla). Dark bars represent performance using the model presented in panel A, light bars represent the “worst” model with large effective population sizes, in the case of hmmix to simulate the data to detect fragments, in the case of *S\** to obtain the expected distribution of *S\** scores. **C** Posterior distributions for the archaic introgression proportion, time of archaic introgression, and gorilla-ghost split time. The dotted line indicates the prior distribution. The black line indicates the posterior inferred with a simple ‘rejection’ algorithm. The red line represents the posterior inferred with neural networks. Compared to the rejection algorithm, neural networks reduce the dimensionality of the summary statistics used and account for possible mismatch between the observed and simulated summary statistics^30^.

### The ghost introgression landscape in eastern gorillas

Having established that a model of ghost introgression into the common eastern ancestor provided the best fit to the empirical data, we aimed to identify the putative introgressed fragments in the genomes of eastern gorillas. To explore this landscape of ghost introgression, we implemented two independent approaches: the *S\** statistic^19,20,31^ and the SkovHMM method, or hmmix^32^. The *S\** statistic detects highly divergent windows relative to an outgroup, under a given demographic model, as introgressed sites^19,20,31^. Hence, the *S\** approach depends on the availability of a demographic model. By contrast, hmmix does not rely on a demographic model to identify putatively introgressed regions, but instead uses the density of private mutations in the ingroup to partition the genome into “internal” and “external” fractions, walking in small windows of 1,000 base pairs (bp) along the genomes^32^. Hence, although both *S\** and hmmix target the same signature of ghost introgression, the algorithms are distinct.

We simulated the expected null distribution of *S\** scores for eastern gorillas with posterior parameter estimates from model A, *i*.*e*. a model without ghost introgression. This yields insights into the presence of any outlier windows in our empirical data using the 99% confidence interval for expected *S\** scores, given the mutation density (number of segregating sites) in each window (Fig. S15, see Methods). Indeed, at this threshold we identify an excess of *S\** outlier windows, suggestive of introgression from an external source into the common ancestor of eastern gorillas: Windows which fall outside the null expectation constitute, on average, 1.64% of eastern lowland genomes and 2.36% of mountain gorilla genomes, respectively (Table S10 - per individual).

We assessed the performance of the *S\** statistic using coalescent simulations where we could trace the introgressed fragments (Methods). The precision and recall are high, with a 90.96% detection rate of true introgressed fragments for eastern lowland gorillas (91.06% for mountain gorillas) at the 99% quantile (Fig 2B, S14, Table S8, see Methods), comparable to the human-Neanderthal scenario^29^. Since the CIs of the null demographic model encompass larger effective population sizes, which would lead to inflated rates of incomplete lineage sorting that might affect the expected distribution of *S\** scores, we also assessed how these parameters influence our findings. Using the maximum values within the 95% CIs, we find that the recall of the *S\** statistic remains high, while the precision falls to 55.82% for eastern lowland gorillas (53.33% for mountain gorillas), reflecting an increase in the false discovery rate, as expected. We conclude that the *S\** statistic performed well in detecting introgressed fragments under our null model, even when assuming misspecification of the null model.

Analogous to previous work^22^, we also employed hmmix to detect introgressed windows^32^, which performs well for the given demographic model (Fig. 2B), with precision and recall well above 80% (Table S9). Considering the strong support for ghost admixture into eastern gorillas, we again used western lowland gorillas as the outgroup and eastern gorillas as the ingroup. We find that 1.48-2.97% of the individual eastern gorilla genomes are inferred as external at a strict threshold for the mean probability of 0.95, with an estimated introgression time of 37-41 kya.

While we observe sharing of the putative introgressed regions across the eastern species, sharing is higher within each subspecies, which again is more pronounced in the mountain than in the eastern lowland gorillas (Fig 3A). This indicates that the majority of the putative introgressed regions are segregating rather than fixed. Pairwise nucleotide differences are elevated between eastern and western gorillas in putative introgressed regions in eastern gorillas, compared to random regions (Fig 3B). Likewise there is an excess of nucleotide differences between individuals of the eastern subspecies in the putative introgressed regions, indicative of an archaic origin of these regions (Fig 3B).

**Fig 3:**
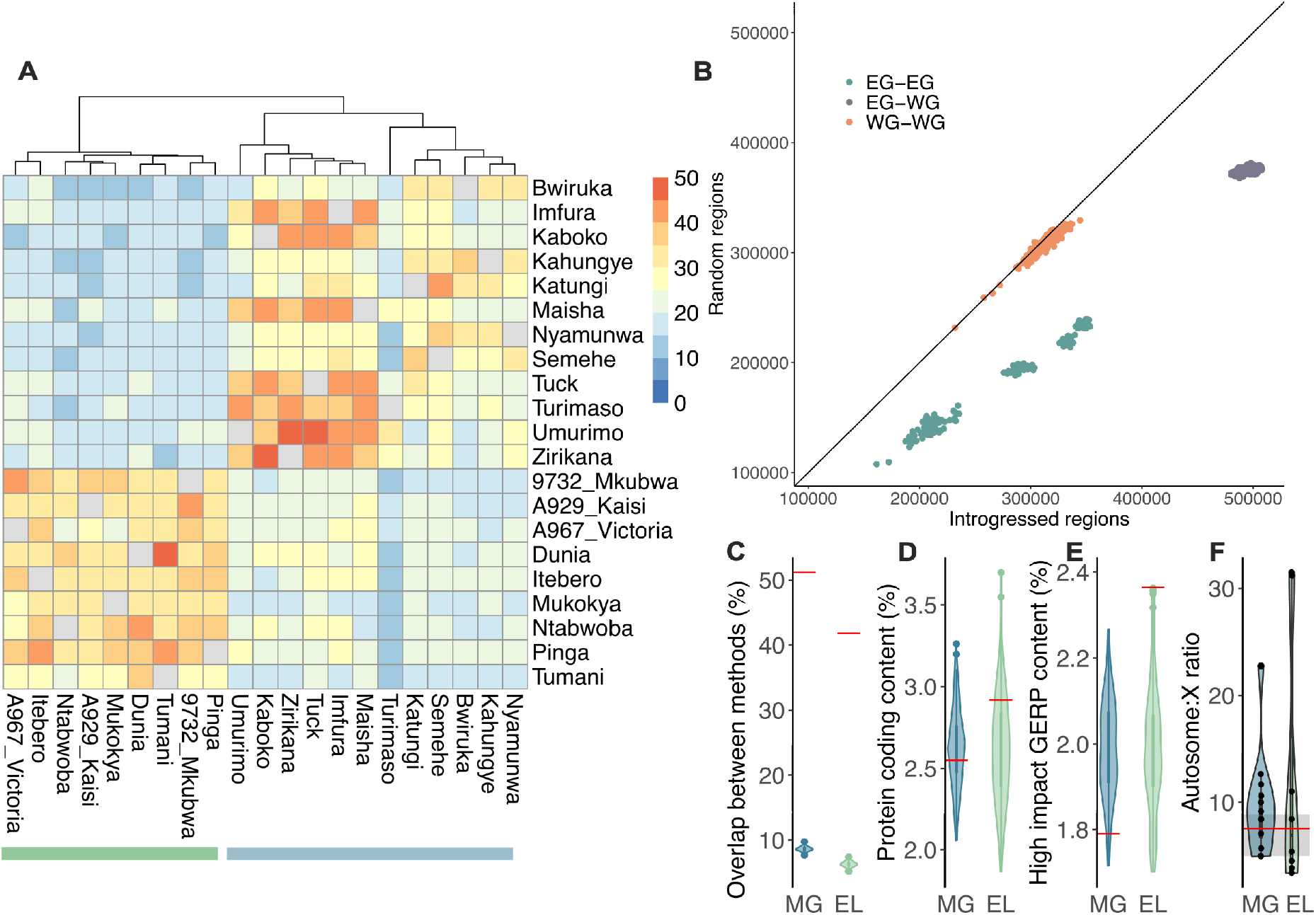
Characterization of introgressed fragments. **A** Sharing of putative introgressed regions across eastern gorillas for autosomal regions detected using the *S\** statistic and hmmix. **B** Pairwise nucleotide differences in introgressed regions (x axis) and in random regions (y axis) matched for length and proportion of positions with sufficient coverage (i.e. avoiding genomic regions without callable sites). Colours indicate the comparison: among eastern gorillas (EG-EG, green), among western gorillas (WG-WG, orange), and between eastern and western gorillas (EG-WG, purple). **C** Percentage of overlapping base pairs in introgressed regions (red lines) and random regions (violin plots) for eastern gorillas. For details on the definition of random regions see Methods. **D** Percentage of protein coding content detected in introgressed regions (red lines) and random regions (violin plots) for eastern gorillas. **E** Percentage of high impact GERP content detected in introgressed regions (red lines) and random regions (violin plots) for eastern gorillas. **F** Autosome:X ratio of introgressed fragments inferred using hmmix for eastern gorillas (violin plots), with reference lines for the equivalent values for bonobos (red line) and humans (distribution as grey bar). In panels C-F, MG = mountain gorillas, EL = eastern lowlands.

The overlap of the autosomal hmmix fragments and the *S\** outliers within each individual is, on average, 42% for eastern lowland gorillas and 51% for mountain gorillas (Table S12). For random genomic regions passing filtering criteria, the observed overlap is, on average, 6% for eastern lowland and 8% for mountain gorillas, suggesting that both methods detect to a large degree the same regions (Fig 3C, Table S15). We thus consider the regions in the intersect of the hmmix outliers and *S\** outliers as our high-confidence putative introgressed regions. The overlap between the two methods increases to 59% for eastern lowland and 68% for mountain gorillas, when using more lenient cutoffs for both methods, *i*.*e*. hmmix fragments of at least 40 kilobase pairs (kbp) and 95% confidence interval *S\** outliers (Table S13). Mountain gorillas (with the exception of Turimaso) consistently exhibit higher proportions of overlapping base pairs of the two methods than the eastern lowland gorillas (Tables S12-S13).

### The interaction of selection and introgression

In contrast to archaic introgressed regions identified in humans and bonobos, the putative introgressed regions in eastern gorillas are not significantly depleted in genic content compared to random genomic regions (Fig 3D). However, we find 127 Mb of autosomal segments longer than 5Mb that are depleted for introgressed fragments (Fig 4). Further, we observe a signal of depletion in archaic fragments on the X chromosome (Fig 3F), on a scale comparable to observations in modern humans^33^ and bonobos^22^. The putative introgressed regions of eastern lowland gorillas exhibit a slightly higher proportion of likely deleterious sites than mountain gorillas, as estimated by the GERP score (Fig 3E). However, under alternate measures of mutational conservation (SIFT, PolyPhen-2 and LINSIGHT scores) the putative introgressed regions of both eastern gorilla subspecies follow random expectation (Fig S23). We also investigated the distribution of gorilla-defined regulatory element annotations from another study^34^. Here, across categories and populations, we only observe an excess of strong enhancers (sE) in mountain gorilla introgressed regions, compared to random regions (Fig S24). These are largely intragenic enhancers (Fig S25), which agrees with patterns of regulatory architecture observed in primate sE^34^.

**Fig 4:**
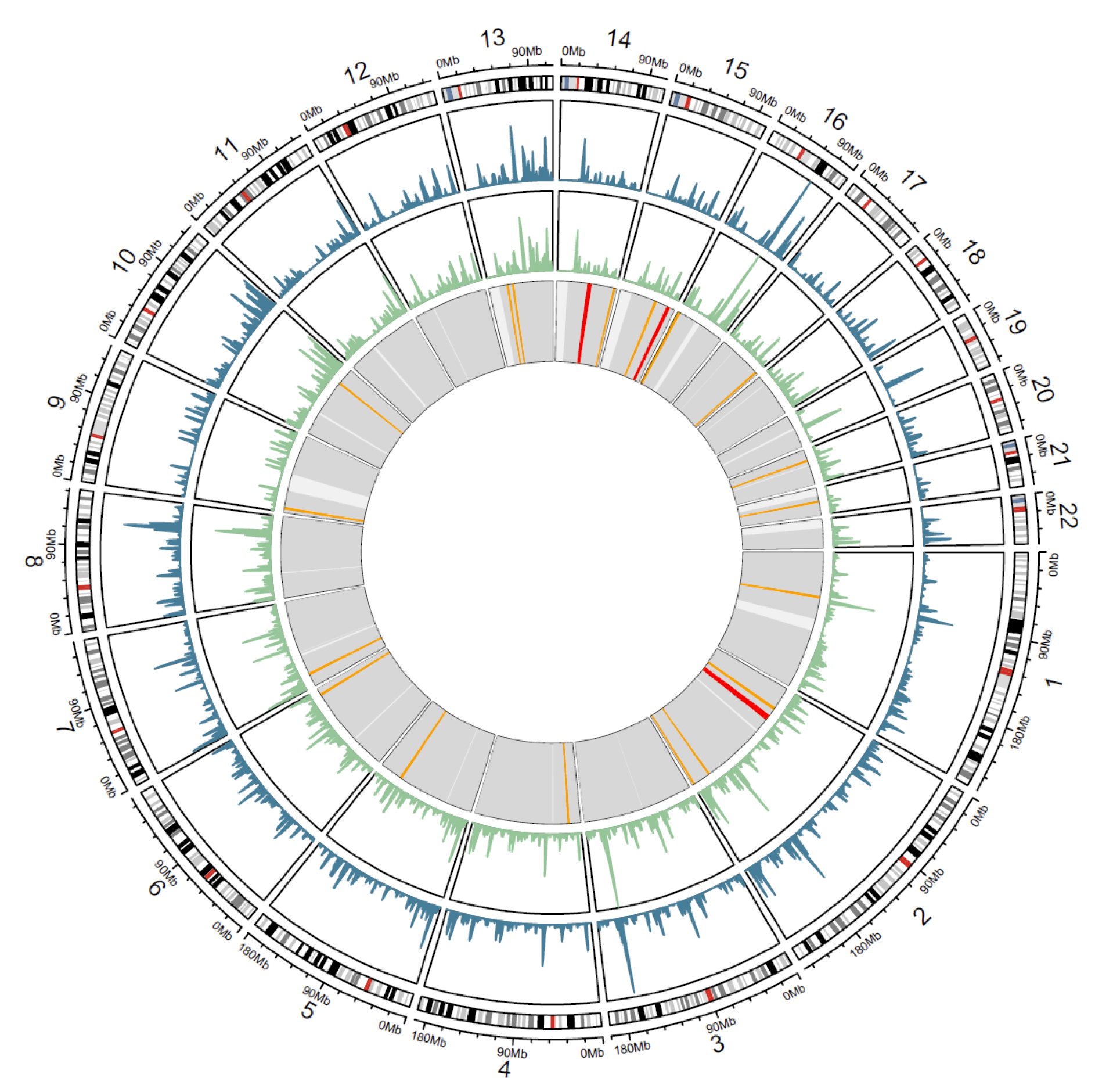
Distribution of introgressed fragments. Outer circle: karyogram of the autosomes based on the human genome (hg19). Second circle from outside: Introgression landscape in mountain gorillas (blue), as cumulative amount of introgressed material in sliding windows of 2 million base pairs (Mbp). Third circle from outside: Introgression landscape in eastern lowland gorillas (green) in sliding windows of 2 Mbp. Inner circle: long regions depleted of introgression content are shown in orange (length >=5Mbp) and red (length >=8Mb). Grey: Genomic regions with sufficient data (>20% of 40 kbp windows passing threshold). White: Genomic regions without sufficient data.

Introgressed fragments can carry beneficial alleles and to explore signatures of adaptive introgression within eastern gorillas, we applied the method VolcanoFinder^35^. VolcanoFinder scans the genome for a signal of a distorted local site frequency spectrum consistent with a selective sweep surrounding an introgressed allele. Outliers of the VolcanoFinder approach (95% composite likelihood ratio) within the putative introgressed regions identified above were considered putative targets of adaptive introgression. We identify seven candidate regions for adaptive introgression (Table S16), three of which are shared between eastern lowland and mountain gorillas. The region with the highest likelihood ratio in VolcanoFinder (chr12:11090005-11324172; max. LR=246.2) contains the bitter taste receptor *TAS2R14*, within which we find several protein-coding changes (Table S17).

## Discussion

Here, we present the first demographic model inferred from representatives of all four extant gorilla subspecies, leveraging the most comprehensive dataset of gorilla genomes available to date, and an improved estimate for gorilla mutation rate from extended pedigree data^36^. The newly sequenced whole genomes of mountain gorillas from Bwindi National Park are genetically close to those from Virunga, but form a distinct cluster within their subspecies (Fig. 1B-C), confirming earlier results from microsatellite data^37^. Eastern lowland gorillas, as represented in our dataset, seem to form a genetically less differentiated population, which includes the individual from Mount Tshiaberimu. Nonetheless, sample size remains a limitation, as high quality invasive samples are highly restricted for endangered species, given ethical and logistical constraints. A more fine-grained analysis of the evolutionary history and population structure of gorillas necessitates denser sampling, which most likely will only be possible through advances in the use of non-invasive samples. For example, a reconstruction of recent patterns of connectivity has been demonstrated from a large panel of faecal samples from chimpanzees^38^. Furthermore, considering the rapid decline of great ape populations over the past centuries, more temporal sampling from historical specimens^24,25^ has the potential to be highly informative on variation lost over time.

Previous estimates of demographic parameters varied greatly under different models, methods and input data^9,13–15^. The ABC approach presented herein leverages population-wise summary statistics. However, since high-coverage, population-level whole genomes are not currently available for Cross River gorillas, a subset of the statistics could not be obtained for this subspecies (Methods, Supplementary Table S6), and those calculated may be relatively less informative (for example, number of segregating sites). For all other populations, multiple individuals were included, yielding a better representation of their diversity in the summary statistics. As such, we have lower confidence in parameters involving Cross River gorillas, such as the relatively large divergence time inferred for the western lowland-Cross River split. This divergence time represents 47% of the inferred eastern-western species split, compared to 26% estimated in a previous study which also inferred a more recent species split time^13^. We note this difference may be attributed to our inclusion of a greater number of western lowlands, known to have high levels of population structure^9,39,40^. We also do not include gene flow between western lowlands and Cross River gorillas as a parameter in our modelling, which would reduce divergence estimates.

The inferred deep divergence time between the two species is at the upper end of previous estimates, and conservative for the detection of putatively introgressed windows under the null model, since larger *S\** scores would be expected to result from an increased number of segregating sites^22^. Indeed, even approximate demographic models with large divergence times may allow a detection of external gene flow into a target population^29^. We demonstrate that the *S\** statistic performed well in detecting introgression under the null model inferred herein (model A), even if the true demography was deviating in terms of ancestral effective population sizes. Demographic modelling presented here finds the best model for gorilla demography to include archaic introgression from an unsampled ‘ghost’ lineage into the common ancestor of eastern gorillas. This accords with a growing literature on the prevalence of introgression from extinct lineages in humans^21,41^, bonobos^22^ and other species^42,43^, as well as theoretical predictions and simulations showing an impact of admixture from unsampled lineages that is likely common rather than exceptional^16,17^. Using extensive simulations, we find strong support for a model including archaic admixture into eastern gorillas, compared to a null model without such ghost admixture, or a model of such an event in western gorillas. The latter may be rather considered similar to a model of deep substructure within gorillas, given the shallower times and small amounts of external gene flow inferred. However, we note that further ghost introgression events may exist beyond what we describe, for example with regards to much smaller amounts of ghost admixture into gorillas, or with shallower divergence times of the ghost lineages, or in the context of larger effective population sizes in western gorillas.

Our inference of 2.47% of ghost introgression is associated with high confidence as the posterior distribution is well differentiated from that of the prior (Fig 2C). This estimate agrees well with the estimates of genome-wide introgression proportions per individual inferred using the *S\** statistic and hmmix (Table S10). We likely underestimate the timing of archaic introgression, since shorter introgressed fragments are more likely to be missed, and another potential complication might be relatively high levels of homozygosity in eastern gorillas^8^, leading to increased haplotype lengths. Our definition of putative introgressed regions as the overlap of outliers inferred with both the *S\** and hmmix methods (Fig 3C) is conservative and on the order expected for these methods, considering their relatively high false-positive rates^29^. Nonetheless, these methods are currently the only reliable tools available for detecting introgressed fragments in comparably small datasets of non-human species, without the availability of a source genome^29^.

A higher degree of sharing of putative introgressed fragments is observed among mountain gorillas than in eastern lowlands (Fig 3A). This is consistent with smaller effective population sizes of these populations, increasing the impact of drift on introgressed genetic variation^18^. High levels of genetic drift and reduced efficacy of natural selection likely also explain the absence of a detectable depletion of genic content in introgressed regions, in contrast to observations in introgressed material of humans and bonobos. Likewise, we do not observe a coherent signature in mutational tolerance in gorilla introgressed material across different metrics, possibly due to genetic drift. Despite this, we do find a number of “introgression deserts”, *i*.*e*. regions depleted of introgressed material in the population (Fig 4), possibly as a result of purifying selection^18^ shortly after the introgression took place. Furthermore, we observe a reduction of introgression on the X chromosome, as also seen in humans and bonobos^22,32,33^. This is most likely a result of strong purifying selection against introgressed variation, as seen in humans and other species^18,32,44^, possibly as a result of a combination with multiple factors^45^. Biased dispersal patterns^46^ and high reproductive skew of gorilla males^47^ might have led to a further reduction of the male-haploid X chromosome in introgressed material. Even though the observed patterns are likely a combination of these factors, we can currently not discern their respective contributions.

We note that our definition of adaptive introgressed targets is highly conservative, as the intersection of the outliers of three different methods *S\**, hmmix and VolcanoFinder as putative adaptive introgressed targets. However, in being conservative we aim to minimise the impact of potential false positives, which is a known caveat of the VolcanoFinder method^35,48^. However, at present this is the only method available to localise signatures of adaptive introgression without a source genome. Interestingly, three candidate genes contain putative functional variants segregating in eastern gorillas and fixed ancestral in western gorillas. One of these genes is *TAS2R14*, which encodes a taste receptor implicated in perception of bitter tastes^49^ and contains six missense variants. Eastern gorillas typically have more herbaceous diets than the frugivorous western gorillas^11^, as such taste receptors are plausible targets of adaptive introgression in eastern gorillas. Bitter taste receptors have been suggested as targets of recent positive selection in western lowland gorillas as well, including a region encompassing *TAS2R14*^13^. It is possible that different mutations in the same region have been under selection in the different species. This could be interpreted in terms of the essential role of taste receptors to avoid toxicity. The gene *SEMA5A* contains a missense variant and a splice region variant; this gene has been associated with neural development, with implications in autism spectrum disorder^50^. However, the functional impact of these variants in gorillas demands further work in the future. Here, we do not find a contribution of adaptive introgression to altitude adaptation, a phenomenon observed in humans and other species^18,51^. In mountain gorillas and eastern lowland gorillas at high altitude, this adaptation is likely driven by different mechanisms, such as the oral microbiome^52^.

In conclusion, our work contributes improved resolution to our understanding of the evolutionary history of eastern gorillas. Across individuals, we recover a putative 16.4% of the autosomal genome of an extinct lineage (Table S14), adding to a growing literature revealing unsampled, now extinct lineages via analysis of variation present in present-day individuals.

## Methods

### Samples and Sequencing

Six eastern gorillas were sequenced as part of this study. Five Bwindi mountain gorillas were sampled after death by the Mountain Gorilla Veterinary Project. One Mount Tshiaberimu individual was sampled under anaesthesia. Convention on the Trade in Endangered Species of Wild Fauna and Flora (CITES) permits were obtained for all samples. Sequencing was performed on the Illumina HiSeq X platform. These six samples are publicly available in the European Nucleotide Archive (ENA) under the project number: PRJEB12821. Detailed information on all samples is provided in the Supplementary Materials (Table S1). Scripts used for data analysis are available on Github under https://github.com/h-pawar/gor_ghost_introg.

### Data processing

We integrated the newly sequenced samples alongside previously published, high-coverage genomic data^7,8^. Raw sequencing reads were mapped to the human hg19 reference genome, as described previously^53^. Given that the hg19 reference does not belong to any of the gorilla subspecies, equal mapping bias will be exerted across all gorillas in our dataset. This would not have been the case if the gorilla reference genome was used instead, as it might have introduced bias in amounts of allele sharing, as observed previously for chimpanzees and bonobos^53^. The final dataset derives from 49 gorillas of known subspecies: 12 mountain (*Gorilla beringei beringei)*, 9 eastern lowland *(Gorilla beringei graueri)*, 1 Cross River *(Gorilla gorilla diehli)* and 27 western lowland *(Gorilla gorilla gorilla)* gorillas.

Processing of data to obtain genotypes followed procedures described in Kuhlwilm et al.^22^. We used bcftools to retain genotypes with a coverage larger than 5-fold and lower than 101-fold, a mapping quality over 20, a proportion of MQ0 reads under 10%, and an allele balance of more than 0.1 at heterozygous positions; bedtools and jvarkit^54^ to filter the data by known repeats (RepeatMasker) and mappability (35 k-mer). Following a previous study^22^, we used the rhesus macaque reference genome (Mmul10) to infer ancestral allele states at each site and generate an ancestral binary genome, as implemented in the freezing-archer repository (https://github.com/bvernot/freezing-archer). Only positions with genotype information in all individuals after filtering were used for calculating summary statistics for the demographic model and the *S\** analysis. For hmmix, missing data was allowed, genotypes were filtered for known repeats and mappability, and then an individual-based filtering was applied for sequencing coverage (depth 6-100), mapping quality (20) and retained only bi-allelic single nucleotide variants (SNVs).

### Demographic modelling

#### Null demographic model

To infer a reliable null demographic model for the four extant gorilla subspecies, we performed Approximate Bayesian Computation (ABC) modelling using the R package abc^30^ with neural networks, following a previously described strategy^22^. Previous demographic models did not include all of the four extant gorilla subspecies^8,13^. We first attempted a merging of these models (Table S7), but in simulations this proved a poor fit to the empirical data in terms of the distributions of segregating sites, one of the main determinants of *S\** (Fig S4).

We used ms^55^ to simulate data, and aimed to generate 35,700 coalescent simulation replicates, of which 35,543 were successful, whereby per iteration we generated 2,500 windows of length 40,000 base pairs (bp), randomly sampling from wide uniform priors informed by the literature^8,13,36^ (Table S3). We sampled local mutation rates from a normal distribution with mean of 1.235e-08 (mutation rate per generation), recombination rate from a negative binomial distribution with mean of 9.40e-09 and gamma of 0.5 and assumed a generation time of 19 years^36^. We scaled the mean mutation rate to 1.976 (1.235e-08 * window size of 40,000 * 4 * Ne of 1000) with a scaled standard deviation of 0.460408 (1.976*0.233). We also scaled mean recombination rate to 1.504 (9.40e-09 * 4 * Ne of 1000 * window size of 40,000). Per window and per population, we calculated the following summary statistics: mean and standard deviations of heterozygosity, nucleotide diversity (pi) and Tajima’s D, as well as the number of population-wise fixed and segregating sites, the number of fixed sites per individual and pairwise *F*_*ST*_ (Table S6). These measures constitute the input summary statistics for all ABC analyses performed in this section. Given only one diploid sample is available for *G. gorilla diehli*, we did not use standard deviations of heterozygosity, nucleotide diversity and fixed sites per individual, as well as mean nucleotide diversity for this population.

We calculated the equivalent summary statistics normalised by data coverage for the empirical data, which had been pre-filtered by repeats, mappability and sufficiently informative windows (>50% of sites with confident genotype calls in all individuals). We also filtered by sites fixed across all gorillas relative to the human reference genome. We accepted parameter values from the prior distribution if they generated summary statistics close to those of the empirical data. This was assessed using a tolerance of 0.005, logit transformation of all parameters and 100 neural networks in the ABC analysis.

#### Alternative demographic models

We performed parameter inference for two further demographic models, in which we allowed gene flow from a ‘ghost’ lineage into the common ancestor of B) eastern gorillas (*G. beringei beringei, G. beringei graueri*) (Table S3) and C) western gorillas (*G. gorilla gorilla, G. gorilla diehli)* (Table S3). For each alternate demographic model, as above, we performed ABC analysis using 35,700 simulation replicates, whereby per iteration we generated 2,500 windows of length 40kbp. We fixed parameters with narrow CIs from model A, in order to reduce the complexity of these models. To assess the impact of fixing well-inferred parameters from the null model on subsequent ghost parameter inference and explore the ghost parameter space more fully we undertook a revised modelling approach (Supplementary Material). In these revised ghost models, we performed parameter inference sampling all parameters from priors, for ghost gene flow into the common ancestor of D) eastern gorillas and E) western gorillas (Table S3). We observed a strong correlation between the estimated parameters of the original and the revised ghost models, albeit with wider posterior distributions for the revised models due to increased complexity and larger parameter space (Supplementary information).

To compare the three main demographic models A) null demography, B) ghost gene flow into the eastern common ancestor, C) ghost gene flow into the western common ancestor, we simulated 10,000 replicates of 250 windows of 40kbp length, fixing the parameters as the weighted median posteriors for each model. To achieve an equal simulated timeframe (number of generations) in all models under comparison, we added a non-interacting ghost population to the null demography, with a divergence time between ghost and extant gorillas equal to that inferred under Model B above. To determine if the models could be differentiated from each other we performed cross-validation with the function cv4postpr (nval=1000, tol=0.05, method=“neuralnet”). We calculated the posterior probabilities of each demographic model using the function postpr (tol=0.05, method=“neuralnet”). The resulting confusion matrix is shown in Table S4. We also performed cross validation and model comparison for the five demographic models: A) null demography, B) ghost gene flow into the eastern common ancestor, C) ghost gene flow into the western common ancestor, D) revised model of ghost gene flow into the eastern common ancestor and E) revised model of ghost gene flow into the western common ancestor, where we still observed model B having the highest support (Table S5, Supplementary Material).

### Detecting introgressed fragments

Following ^20,22,31^ we calculated the *S\** statistic using a customized version of the package freezing-archer, accommodating non-human samples. We calculated the *S\** statistic genome-wide in 40kbp windows, sliding every 30kbp, using the following test (i = ingroup) and reference (o = outgroup) populations: 1) GBG (*G. beringei graueri*-i, *G. gorilla gorilla*-o) and 2) GBB (*G. beringei beringei*-i, *G. gorilla gorilla*-o). For the *S\** analysis, 15,181,832 variants were included.

Identifying outliers for the *S\** statistic requires a distribution of scores for local mutation densities (represented by numbers of segregating sites in the dataset) under a demographic scenario without introgression, as the null model. We used the weighted median posteriors for each parameter value from the above ABC analysis to generate simulated data, specifying the number of segregating sites in a stepwise manner (from 15 to 800 in steps of 5). For each stepwise segregating site (158 in total), we simulated 20,000 windows of length 40kbp, to which we applied the *S\** statistic for each of the scenarios (GBG, GBB). From this we obtained generalised additive models (GAMs) per scenario for three confidence intervals (95%, 99%, 99.5%) using the R package mgcv, following the procedures described in detail in ^22,31^. From these GAMs, we predicted the expected *S\** distributions under the null model without archaic introgression. Applying the GAMs to the empirical data we inferred whether any windows lay outside the expectation per scenario and per confidence interval, assessing confidence intervals of 95%, 99% and 99.5%. As such, the threshold of significance is defined as the 95%, 99% or 99.5% confidence interval from the standard deviation for expected *S\** scores, given the mutation density^22,31^.

To assess the performance of the *S\** statistic under our null model and its robustness to model misspecifications, we performed validation analyses following Huang et al.^29^, using msprime^56,57^ simulations with explicit tracking of the introgressed fragments. Briefly, we simulated expected distributions of *S\** scores for the null model (model A), and for a model where the effective population sizes before 40 kya were set to the upper end of the 95% CI (“worst” null model, in terms of highest expected amount of incomplete lineage sorting). We then simulated datasets of 10 outgroup individuals (western lowland gorillas) and a single ingroup individual (eastern lowland or mountain gorillas) for model B and a model where the effective population sizes before 40 kya were set to the upper end of the 95% CI (“worst” model B). We then obtained putatively introgressed fragments using the expected scores from either model A or the “worst” null model (Table S8). For each model, we performed 100 replicates and calculated the average precision and recall at different thresholds. For the *S*\* approach, we used the quantiles of the *S*\* statistic as thresholds, which range from 0 to 0.999.

In an independent approach to the *S\** statistic, we applied hmmix^32^. We obtained the input files for this method: weight files, local mutation rates and individual observations files using scripts provided with the repository for hmmix (https://github.com/LauritsSkov/Introgression-detection, as of 2018/08/02), as well as bcftools, bedtools, jvarkit and custom R scripts. The macaque allele (RheMac10 assembly) was used for polarisation of alleles. We then applied the method to the eastern gorillas using the following prior parameters: starting_probabilities = [0.98, 0.02], transitions = [[0.9995, 0.0005], [0.012, 0.988]], emissions = [0.05, 0.5]. We confirmed that using different parameters did not affect the results. We used a recombination rate of 9.40× 10^−9^ per site per generation and 19 years generation time with the median fragment length to estimate introgression time. Decoding, *i*.*e*. assigning internal and external states to specific genomic regions, was done with the script provided with the repository. Putative external fragments were filtered for posterior probabilities of 0.9 (lenient) or 0.95 (strict), and required to contain at least 5 private positions. We also conducted a performance analysis of hmmix on introgressed fragments in simulations of either model B or the “worst” model B, with results similar to those for *S*\* (Table S9). For performance testing the hmmix approach, we used the posterior probabilities estimated by hmmix as thresholds, which range from 0 to 0.9999.

We note that only hmmix could be used to infer archaic introgressed fragments on the X chromosome, due to the lack of a gorilla demographic model for the sex chromosomes.

### Exploring introgressed regions

To obtain a consensus set of putative introgressed regions, we overlapped the autosomal outlier regions inferred under the two methods within each eastern gorilla. For this overlap, we calculated the percentage of overlapping base pairs, considering in turn each *S\** confidence interval (95%, 99%, 99.5%), and with and without a 40kbp length cutoff for hmmix regions identified under the strict threshold. Imposing a 40kbp length cutoff retains 76.7% of the total strict hmmix regions (Tables S10,S12). We consider the intersect of the *S\** 99% outliers with the strict hmmix autosomal outliers, as our putative introgressed regions of high confidence. To determine whether the overlap obtained differed from random expectation we generated intersections of random regions, of equivalent distribution to the empirical data, for 100 iterations.

As a proxy for gene density we calculated the proportion of protein coding base pairs within these regions of high confidence. As above, we compared this to the proportion of protein coding base pairs within 100 iterations of random genomic regions, of equal length distribution as the putative introgressed regions within each eastern gorilla. We calculated pairwise nucleotide differences between individuals in the putative introgressed regions and in random genomic regions of equal length distribution and sufficient callable sites. This was conducted for three comparisons 1) among eastern gorillas 2) among western gorillas 3) between eastern and western gorillas.

To assess mutational tolerance, we used GERP, SIFT, PolyPhen-2 and LINSIGHT scores^58–61^. We calculated the proportion of high impact sites for GERP, SIFT and PolyPhen-2 scores and the mean LINSIGHT score within putative introgressed regions and random regions of equal length distribution and sufficient callable sites. To explore the impact of introgression on regulatory elements, we calculated the proportion of regulatory base pairs using gorilla-defined regulatory element annotations^34^, within putative introgressed and random regions of equivalent length and callability. This was assessed globally and per regulatory element type, considering poised, strong and weak, enhancers and promoters.

We further explored our putative introgressed regions of high confidence using PCA (Fig S17). This was generated using the biallelic sites in our putative introgressed regions. For comparison, we also generated PCAs of one random set of random regions, with equal length distribution of random regions as the putative introgressed regions per eastern gorilla. The PCAs in Fig 1 were generated using biallelic SNPs of random genomic regions of equivalent length distribution to the putative introgressed regions of GBB Bwiruka. This sample of random genomic regions is representative of the whole genome. All PCAs were generated with the R package adegenet^62^. We generated phylogenetic trees of our putative introgressed regions and one random replicate (Fig S18), using the ‘K80’ model of nucleotide substitution, using the adegenet package^63^. Haplotype networks were drawn using pegas^64^.

We localised introgression deserts by screening 1Mb non-overlapping windows (‘bins’) spanning the genome. We filtered out bins overlapping centromeres and those at the end of each chromosome which were < 1Mb in size. Per bin we calculated the frequency of putative introgressed regions falling within the bin, for each eastern gorilla. We also calculated data coverage of the bins and filtered by mean callable proportion > 0.5. Deserts hence constitute bins where no eastern gorilla carried a putative introgressed region, and which had a reasonable number of callable sites.

Plots were created with ggplot2^65^, circlize^66^ and pheatmap (https://github.com/raivokolde/pheatmap). Genomic ranges were analysed with the GenomicRanges package^67^.

### Adaptive introgression

To explore signatures of adaptive introgression within eastern gorillas, we applied the genome-wide scan VolcanoFinder^35^. To do so, we polarised the data to two outgroups. First, we polarised the human reference allele using the rhesus macaque allele, and subsequently polarised the gorilla genotypes by this polarised allele representing the ancestral state. To obtain the allele frequency input files per chromosome, we then filtered our data to only eastern gorilla genotypes at biallelic sites, and also filtered out sites with multiple ancestral alleles (where polarisation would be uncertain) and sites of reference homozygotes. The second input file required is an empirical unnormalised site frequency spectrum (SFS), which we generated by obtaining the unfolded SFS, normalising so all site categories sum to 1 and then filtering out the first category (the 0 entry). We called VolcanoFinder specifying “-big 1000, D = -1, P = 1, Model = 1”. For computational efficiency, we performed the VolcanoFinder scan in blocks, whereby each chromosome was split into blocks of approximately equal numbers of base pairs. We placed a test site every 1,000 bp (-big 1000). We set D to -1, so VolcanoFinder iteratively tested a grid of values for genetic distance internally and selected the value that maximises the likelihood ratio^35^. We set P to 1 as our input data was polarised. We used Model = 1, following procedures applied to human data^35^, as well as non-human species^68,69^.

We took the 95% outliers of composite likelihood ratio (CLR) scores calculated from VolcanoFinder and intersected these regions with our putative introgressed regions (identified above), to obtain putative adaptive introgressed targets. To explore potential functional consequences, we assessed which genes and which mutations fall within the putative adaptive introgressed targets, using the Variant Effect Predictor annotation (Version 83)^70^.

## Acknowledgments

We thank Derek Setter for valuable guidance in applying VolcanoFinder. We thank the Uganda Wildlife Authority for the Gorilla monitoring and research permission. We are grateful to the Life Science Compute Cluster (LiSC) of the University of Vienna. This project has been funded by the Vienna Science and Technology Fund (WWTF) [10.47379/VRG20001] to M.K.; the European Research Council (ERC) under the European Union’s Horizon 2020 research and innovation programme (grant agreement no. 864203), PID2021-126004NB-100 (MINECO/FEDER, UE), Secretaria d’Universitats i Recerca and CERCA Program del Departament d’Economia i Coneixement de la Generalitat de Catalunya (GRC 2021 SGR 00177) to T.M.-B.; H.P. was supported by a Formació de Personal Investigador fellowship from Generalitat de Catalunya (FI_B100131); M.A.-E. was supported by an FPI (Formación de Personal Investigador) PRE2018-083966 from Ministerio de Ciencia, Universidades e Investigación; C.T.-S. and Y.X. were funded by the Wellcome grant 098051; K.G. was supported by the Swedish Research Council grant (2020-03398); María de Maeztu Mobility Fellowship to J.L.K; O.D. was supported by a John Templeton Foundation grant (ID: 62178); A.A. received funding from UCL’s Wellcome Trust ISSF3 award (204841/Z/16/Z); Q.A. is supported by strategic funding from Monash University (STG-000114).

