## Supplementary text and figures for "Ghost admixture in eastern gorillas"

**Supplementary Material**

**1. New data generation and quality assessment	2**

**2. Exploratory phylogenomic analyses	3**

**3. ABC modelling	7**

3.1 Initial demographic modelling based on the literature	7

3.2 Summary statistic decorrelation	9

3.3 Adjusted demographic modelling	10

3.4 Revised demographic modelling	16

3.5 Parameters from hmmix	21

3.6 Validation of method performance	21

**4. Characterising introgressed fragments	23**

4.1 General features	23

4.2 Introgression and selection	27

4.3 Functional consequences: mutational tolerance	29

4.4 Functional consequences: regulatory elements	31

**References	35**

### 1. New data generation and quality assessment

Six mountain gorilla samples from the Bwindi national park were obtained from deceased individuals as part of the Mountain Gorilla Veterinary Project. Katungi and Kahungye died as infants. Semehe, Nyamunwa, Nkuhene and Bwiruka died as adults. A male Mount Tshiaberimu individual, Mukokya, was sampled under anaesthetic, an intervention as part of a study on the long-term survivability of the small group of Mount Tshiaberimu gorillas (numbering six at the time of sampling). All these samples were imported into the UK in compliance with the legislation for endangered species (CITES).

DNA was extracted from these samples and sequenced on Illumina Hiseq X to 90Gb per sample using non-PCR libraries. After mapping, we performed quality controls and dropped the sample Nkuhene due to very low quality, with 80% of read duplicates and 2X average coverage. The rest of the samples performed similarly to previous mountain gorilla and eastern lowland gorilla samples included in this study.

### 2. Exploratory phylogenomic analyses

Numerous possible ghost introgression scenarios exist in the context of a two clade topology, such as that of the gorillas. To explore the demographic history of gorillas we performed initial exploratory phylogenomic analyses, f-statistics and the admixturegraphs method as implemented in admixtools2 (Maier et al. 2023). Briefly, we converted the genotypes of the autosomes after quality filtering (Methods) to the eigenstrat format, adding one *Pongo pygmaeus* individual (SRS396836) as an outgroup (Prado-Martinez et al. 2013), and retaining only positions where more than 25 individuals had high-quality genotypes. We then calculated pairwise f2-statistics (blgsize=500000) for the four gorilla subspecies and the orangutan individual. Then, we used the find_graphs function to determine the best fitting graphs with an increasing number of admixture edges from 0 to 5 (Fig. S1), defining the orangutan individual as outgroup. The best graph without admixture correctly separates the two gorilla species. The best graph with one admixture edge likely represents substructure in western lowland gorillas, although with an admixture proportion of 0%. Still, this graph fits significantly better (bootstrap p-value 0.0002) than the graph without admixture edges. Further edges increase complexity first in western, then also eastern gorillas, but do not significantly differ from less complex graphs (bootstrap p-value >0.05). We caution that with increasing complexity and in the absence of a hypothesis, the reliability of this method is limited, as discussed extensively by the authors of the method (Maier et al. 2023). Furthermore, with the large space of possible graphs when involving many recent and ancestral populations, different graphs are inferred when repeating the inference (Sorensen *et al.*, *in press*). We also explicitly tested a graph with ghost admixture into the ancestor of eastern gorillas (Fig. S2). This graph provides a better fit than one without admixture edges (p=0.002), but worse than the best graph with one edge (p=0.002).

A more general constraint is that if there had been ghost admixture into any of the four terminal populations (mountain gorillas, eastern lowland gorillas, western lowlands gorillas, Cross River gorillas), these statistics could be informative, as asymmetries between the clades would be introduced. Still, these could be confounded by gene flow between the terminal clades. When explicitly testing such asymmetries D(EG, EG; WG, Orang) or D(WG,WG;EG,Orang), we find no such signature for the eastern gorilla populations, and a weak signature (z score <4) of allele sharing between either Cross River gorillas and eastern gorillas or western lowland gorillas and the outgroup, as shown below (Table SA).

| pop1 | pop2 | pop3 | pop4 | f4 score | z |
| --- | --- | --- | --- | --- | --- |
| MG | ELG | WLG | Orang | 0.000002 | 0.07 |
| MG | ELG | CRG | Orang | -0.00004 | -1.5 |
| WLG | CRG | MG | Orang | -0.00008 | -2.33 |
| WLG | CRG | ELG | Orang | -0.0001 | -3.67 |

Table SA: f4-statistics of the population configuration f4(pop1,pop2;pop3,pop4), with corresponding z-scores.

As such, we focus on exploring possible ghost introgression events into the common ancestor of either eastern or western gorillas. These represent biologically plausible scenarios, which f4-statistics and the admixture graphs method are not able to detect. Instead, we can apply statistical methods developed to detect introgressed fragments in individual genomes from an unsampled or ‘ghost’ population (*S** and hmmix), see Methods. We note that these methods require an ingroup population, which experienced introgression and an outgroup population, which did not. Hence a scenario of ghost introgression into the common ancestor of all extant gorillas would be undetectable under current approaches.

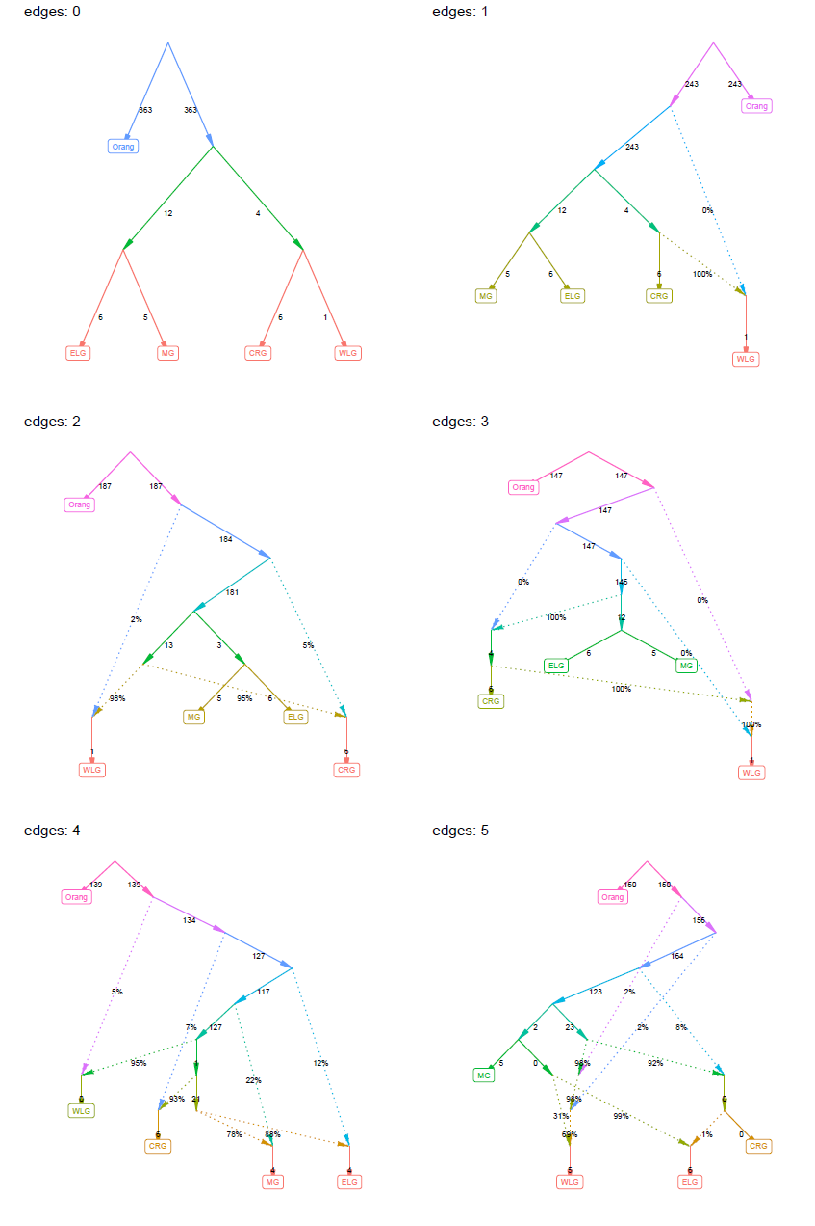

**Figure S1:** Best fitting admixture graphs for up to five admixture edges for the four gorilla subspecies.

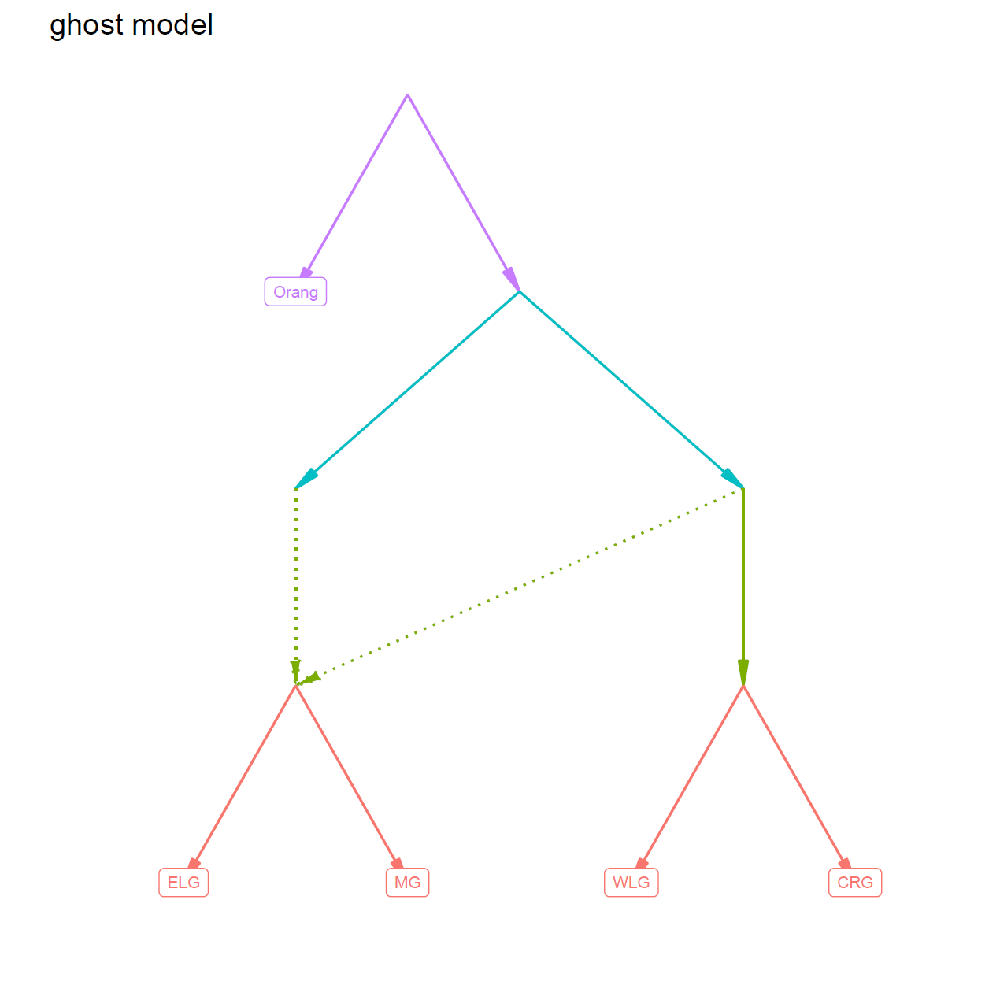

**Figure S2:** Admixture graph with a ghost population contributing to the ancestor of eastern gorillas.

### 3. ABC modelling

The workflow of the main analyses regarding demographic modelling and putative introgressed fragments is shown in Fig. S3.

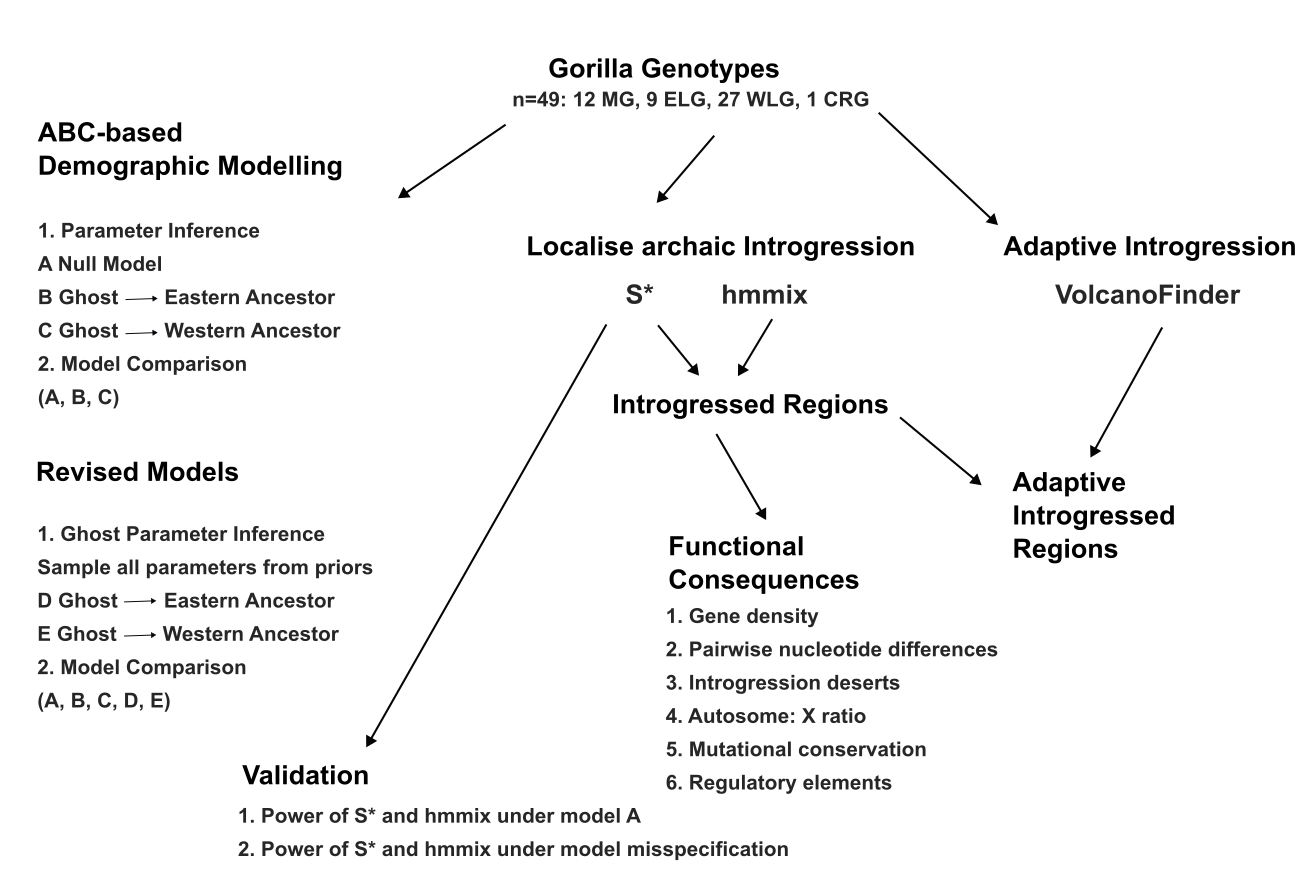

**Figure S3:** Workflow of the main analyses.

#### 3.1 Initial demographic modelling based on the literature

Exploring the question of archaic introgression with the *S** statistic (Plagnol and Wall 2006; Vernot and Akey 2014) requires a window-based demographic model, to test for outliers of the statistic. However, no previous demographic model had yet included all four known subspecies of gorilla. Moreover, due to the use of disparate data and methodologies previous demographic analyses of gorillas had resulted in widely divergent estimates for key parameters, including the estimated divergence time of the eastern and western gorilla species (Thalmann et al. 2007; McManus et al. 2015; Mailund et al. 2012; Scally et al. 2012).

In an initial approach, we merged the parameters from the two most recent studies estimating gorilla demographic parameters, namely McManus et al. (McManus et al. 2015) and Xue et al. (Xue et al. 2015), see Table S7. McManus et al. (McManus et al. 2015) applied a G-PhoCS approach to estimate current and ancestral population sizes, divergence times and gene flow between 9 western lowlands, 2 eastern lowlands and 1 Cross River gorilla in the model. We used parameters estimated by McManus et al. (McManus et al. 2015) under a human-gorilla divergence time of 12 mya, since this was the closest to the 13 mya human-gorilla divergence time more recently inferred by Besenbacher et al. (Besenbacher et al. 2019). Xue et al. (Xue et al. 2015) newly sequenced mountain gorillas from the Virunga subpopulation and inferred effective population sizes and divergence times from PSMC analysis.

We simulated this merged model in msprime (Kelleher, Etheridge, and McVean 2016) then in ms (Hudson 2002) in order to sample the mutation and recombination rates from a normal distribution and a negative binomial distribution respectively. We note that for the gorilla species split time we simulated using a ‘low divergence’ value of 261,000 years ago from McManus et al. (McManus et al. 2015), and a ‘high divergence’ value of 429,000 years ago estimated by Scally et al. (Scally et al. 2012) which is consistent with Mailund et al. (Mailund et al. 2012). The resulting distributions of segregating sites under both simulated models deviated substantially from those obtained using the empirical data (Fig. S4). Under both models, we observe a similar, but larger number of segregating sites compared to the empirical data. As such we embarked on inferring a novel population-level demographic model for the extant gorillas using an ABC approach, as detailed in Methods. We note that both the McManus et al. (McManus et al. 2015) and Scally et al. (Scally et al. 2012) values for the gorilla species split time are substantially lower than that inferred in our ABC parameter inference at a weighted median posterior value of 965,481 years ago. Our estimate of a gorilla species divergence time is within the range of previous estimates, but at the upper end, for example Thalmann et al. similarly inferred a range of 0.9-1.6 mya (Thalmann et al. 2007). Conceptually, a large divergence time is conservative for applying the *S** statistic, as the expected values increase with divergence time (Kuhlwilm et al., 2019). As such, we are conservative in our detection of putative introgressed fragments.

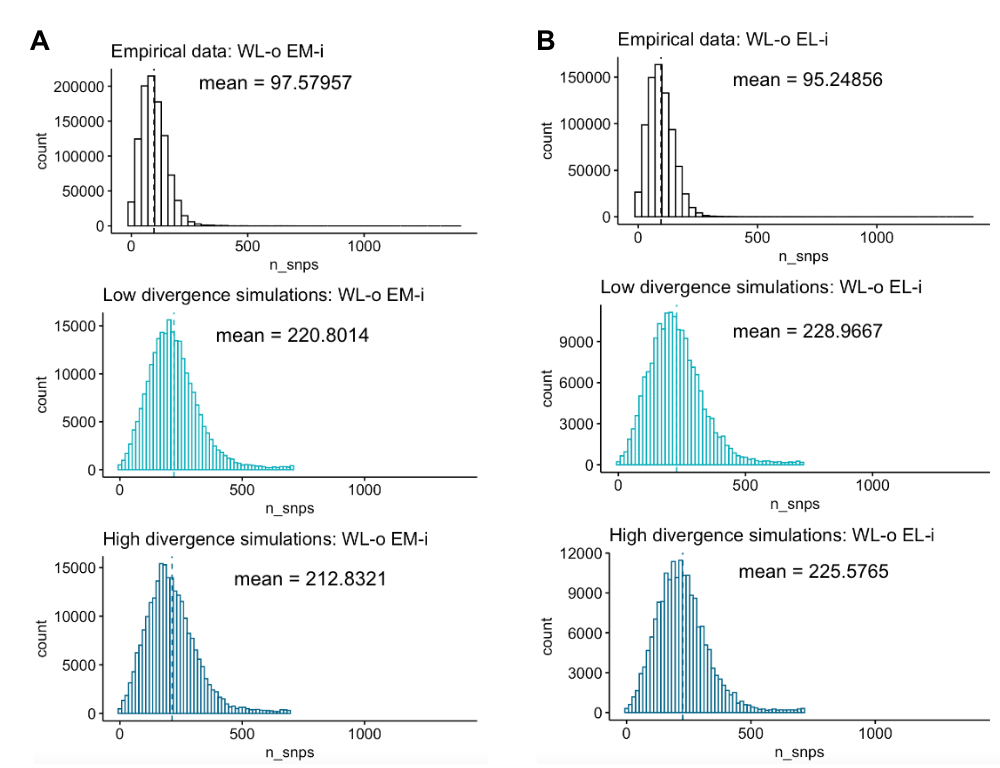

**Figure S4:** Segregating site distributions in the empirical data, low divergence simulations and high divergence simulations under an initial merged model for **A** mountain gorillas and for **B** eastern lowland gorillas.

#### 3.2 Summary statistic decorrelation

We assessed possible correlations between the summary statistics used in the ABC analysis. This could arise, as many of the summary statistics incorporated are related to the SFS. As such, we see substantial correlations between the highly related measures of fixed sites per individual, population-wise fixed sites and population-wise segregating sites (Fig S5A). Correlated summary statistics in the ABC analysis could have two outcomes, it could introduce bias in the posteriors, or alternately, the redundancy simply captures the same information at the expense of adding additional statistics. In principle the neural network method we use to perform the ABC should be robust to any such correlations. Nonetheless we explored the impact of decorrelating the summary statistics on the posteriors obtained.

First, we simplified the correlated summary statistics using an *ad hoc* approach. We summed the correlated statistics (mean and standard deviations of fixed sites per individual, population-wise fixed sites and population-wise segregating sites), and used this one value as an input statistic, alongside the non-correlated statistics. Using this set of ‘ad hoc’ decorrelated summary statistics we performed the ABC analysis of parameter inference for the null model. The resulting posteriors did not differ greatly from those arising when using the correlated summary statistics as input.

Next we performed a formal decorrelation of the summary statistics. Wegmann et al. (Wegmann, Leuenberger, and Excoffier 2009) recommend a partial least-squares (PLS) approach to obtain uncorrelated summary statistics. PLS aims to maximise the covariance between summary statistics and parameters in an approach which is conceptually similar to principal components analysis. Following (Wegmann, Leuenberger, and Excoffier 2009) we applied a Box-Cox transformation on each summary statistic separately, to transform the data to be normally distributed. To perform the Box-Cox transformation and PLS analysis we followed the procedure detailed in the findPLS.R script provided by the ABCtoolbox package (Wegmann et al. 2010). We performed a first pass of the PLS analysis defining 36 components, equal to the number of retained summary statistics after applying the Box-Cox transformation. The resulting root mean square error of prediction (RMSEP) plots indicated that the optimum number of PLS components was 10, which explained 95.6% of the variance (Fig. S5B). As a confirmatory step we re-performed the PLS analysis with the optimum number of components (10) (Fig S5C), following (Luqman et al. 2021). We then performed ABC parameter inference for the null model, using the 10 optimal PLS components as input for the summary statistics. We also performed ABC parameter inference for the subsequently 17 PLS components which explain 98.9% of the variance. In both cases the weighted median posteriors obtained were similar to those obtained using the original summary statistics (which include correlations). As such, we proceed with the original set of summary statistics for ABC analysis in the main text, but we introduce a logit transformation to ensure the posteriors would be within the distribution of the priors (Fig S5D).

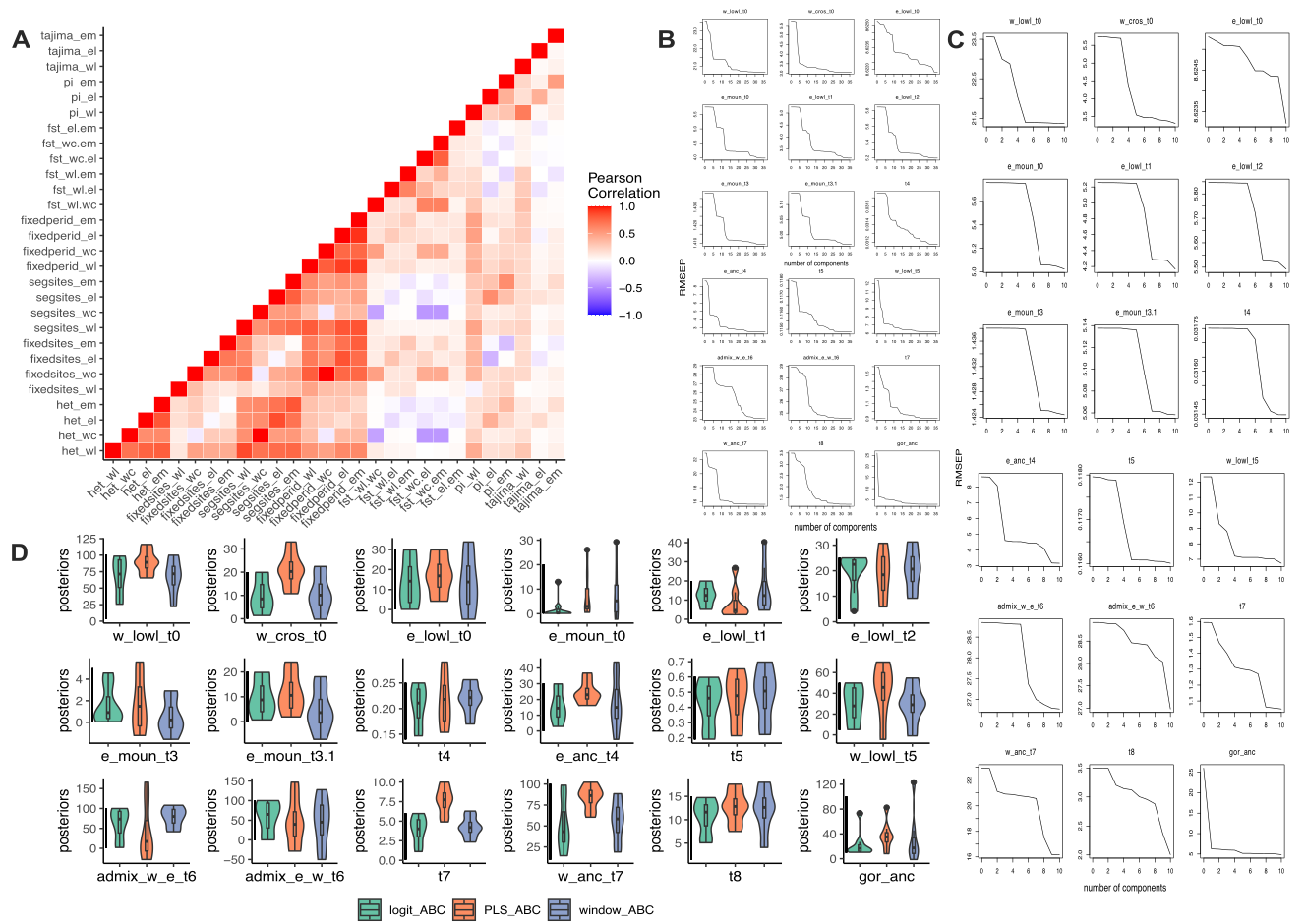

**Figure S5: A** Summary statistic correlations. RMSEP plots with **B** 36 PLS components and with **C** the optimal 10 PLS components. **D** Posterior distributions under the final ABC protocol (teal, ‘logit-ABC’), under the PLS-ABC protocol (red) which takes the 10 PLS components rather than the summary statistics as input and under the initial ABC protocol (purple) which used correlated summary statistics without a logit transformation. The black vertical line represents the prior distribution for each parameter.

#### 3.3 Adjusted demographic modelling

We inferred demographic parameters under a model without admixture from an unsampled lineage (Fig. S6A), as well as a model with such an admixture event into the ancestral western gorilla population (Fig. S6B), as described in Methods. We fixed parameters which were inferred well after inspecting the posterior distributions of the null model (with only the extant gorilla lineages) (Fig. S7, Table S3), and inferred a set of parameters including ghost admixture into the common ancestor of eastern (Fig. S8, Table S3) or western (Fig. S9, Table S3) gorillas, respectively.

In the null model (without admixture from an unsampled lineage) we fix the parameters t1-t3 at the midpoint of their prior ranges, since these are very recent events, with narrow priors. In initial iterations of ABC-based modelling, we observed that parameters t1-t3 were contributing noise, but would contribute little information to the question of deeper demographic history, which is the main focus of the current study.

We also fix the parameter t6 (time of extant admixture between western lowland gorillas and the common eastern ancestor) at 34 kya. This was the result of converting continuous migration implemented by (McManus et al. 2015) to defined migration pusles, using the midpoint between the western subspecies split time inferred by (McManus et al. 2015) and the present. All other parameters were allowed to vary in the null model, sampling from priors informed by previous literature (as detailed in Methods).

We provide a yaml file in the demes format (Gower et al. 2022), as drawn in Fig. S10, for the best supported demographic model for gorillas, which includes a component of ghost admixture into the common ancestor of eastern gorillas.

We explored the impact of fixing well-inferred parameters from the null model on subsequent parameter inference in the ghost models in the section 3.4 Revised simulation approach.

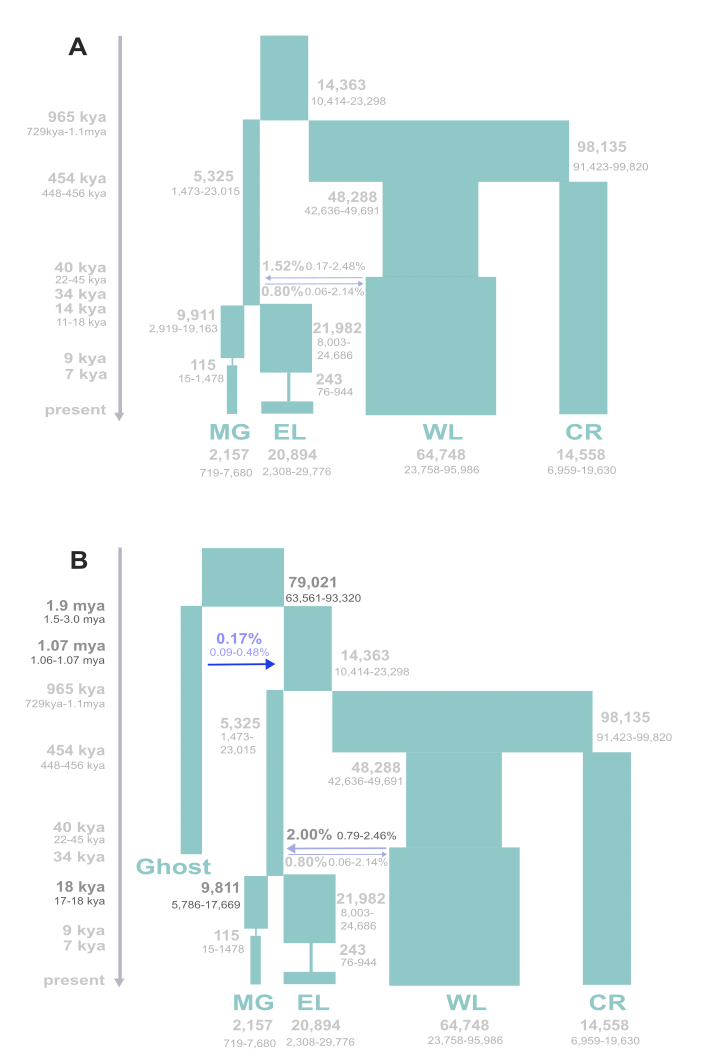

**Figure S6: A** Null model of gorilla population history (only extant populations). 95% credible intervals are shown for all parameters inferred. **B** Alternate model allowing the possibility of ghost introgression into the common ancestor of western gorillas, resulted in a model of ancestral population structure being inferred. We note under a model of ghost gene flow to the western common ancestor, the posteriors indicate a small contribution to the common ancestor of all gorillas (consistent with ancestral substructure), rather than a defined pulse to the western common ancestor. In darker colours are the parameters inferred under this alternate model with their 95% credible intervals.

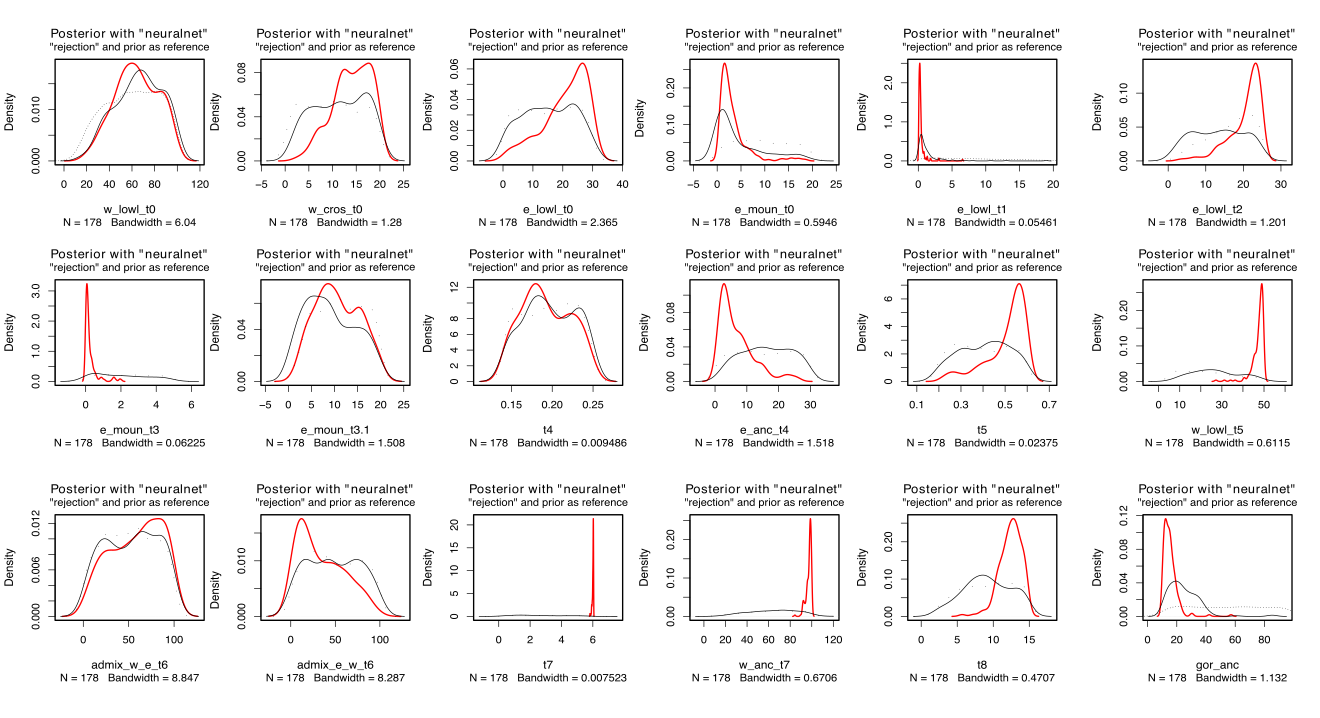

**Figure S7:** Parameter distributions for all parameters inferred under the ABC null model. Red indicates the posterior distribution inferred with neural networks. Black indicates the posterior distribution inferred under a rejection method. The dotted grey line indicates the prior distribution.

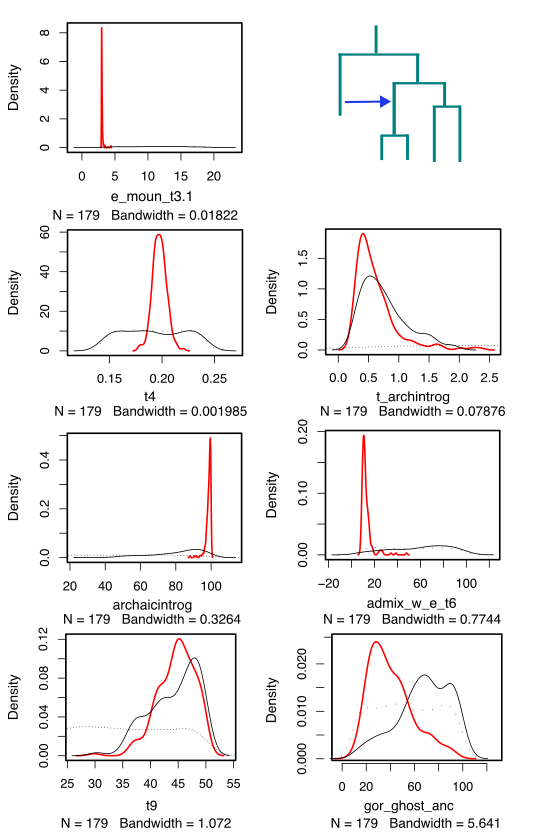

**Figure S8:** Parameter distributions for all parameters inferred under the ABC model allowing gene flow from a ghost lineage into the common ancestor of eastern gorillas. Red indicates the posterior distribution inferred with neural networks. Black indicates the posterior distribution inferred under a rejection method. The dotted grey line indicates the prior distribution.

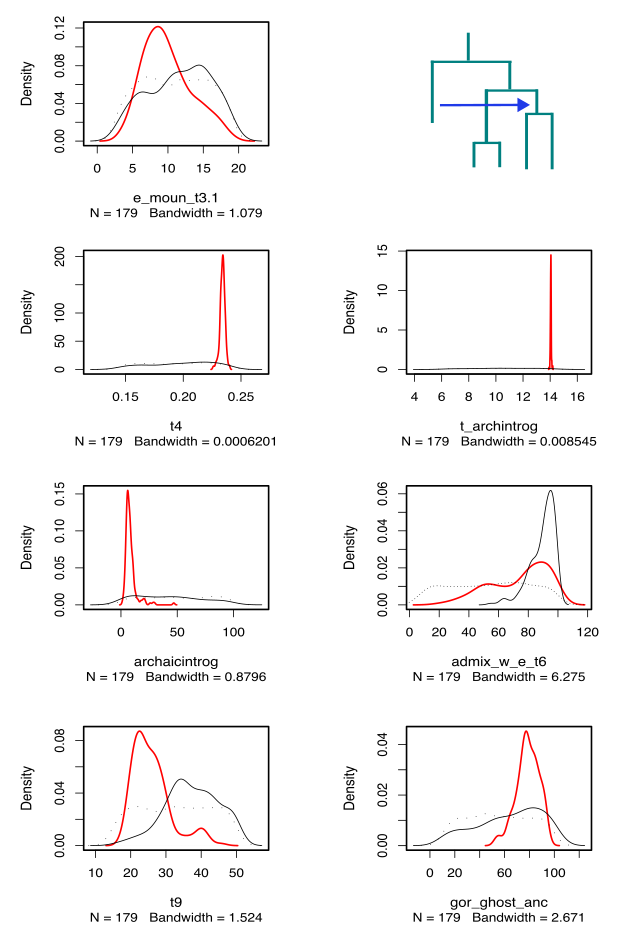

**Figure S9:** Parameter distributions for all parameters inferred under the ABC model allowing gene flow from a ghost lineage into the common ancestor of western gorillas. Red indicates the posterior distribution inferred with neural networks. Black indicates the posterior distribution inferred under a rejection method. The dotted grey line indicates the prior distribution. We note under a model of ghost gene flow to the western common ancestor, the posteriors indicate a small contribution to the common ancestor of all gorillas (consistent with ancestral substructure), rather than a defined pulse to the western common ancestor.

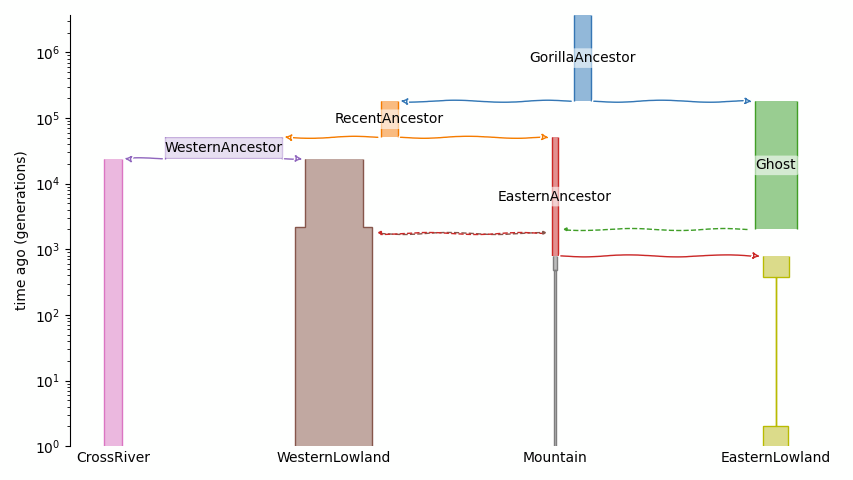

**Figure S10:** Final demographic model with ghost admixture into eastern gorillas, implemented in demes (times in log-scale). The Python package demesdraw was used to generate this figure (https://github.com/grahamgower/demesdraw).

#### 3.4 Revised demographic modelling

In our original ghost models, we fixed parameters with narrow CIs under the null model, in order to reduce the complexity of these models. To explore the ghost parameter space more fully we undertook a revised demographic inference approach for the ghost models, in which we sampled all parameters from priors (Table S3). Again we allowed gene flow from a ghost lineage into the common ancestor of eastern gorillas or western gorillas respectively.

We note that sampling all parameters from priors considerably increases model complexity. Nonetheless, we obtain largely coherent results with those of our original ghost models (in which we fixed parameters well inferred under the null model), albeit with wider confidence intervals inferred (Table S3, Fig. S11-13).

Under the revised modelling, we infer 1.93% of ghost gene flow into the common ancestor of eastern gorillas, (0.33-2.45%, 95% CI) from a ghost population which diverged from extant gorillas ~3.1 Mya (1.43-3.77 Mya, 95% CI). We estimate the timing of ghost gene flow to have occurred 819 kya (146 kya-1.07Mya).

Whereas, for the revised model of ghost gene flow to the ancestral western population, the posterior distribution for the proportion of ghost gene flow tends to 0 (weighted median=0.43%; 0.01-1.98%, 95% CI), which is expected where there is no clear signal of introgression. Moreover, the timing of introgression in this scenario (weighted median=462 kya; 457-472 kya, 95% CI) tends towards the estimate for the extant gorilla species divergence time (weighted median=466 kya; 295-864 kya, 95% CI).

To compare the five demographic models A) null demography, B) original model of ghost gene flow into the eastern common ancestor, C) original model of ghost gene flow into the western common ancestor, D) revised model of ghost gene flow into the eastern common ancestor and E) revised model of ghost gene flow into the western common ancestor, we simulated 10,000 replicates of 250 windows of 40kbp length, fixing the parameters as the weighted median posteriors for each model. We calculated the posterior probabilities of each demographic model using the function postpr (tol=0.1, method="neuralnet"). Model B was overwhelmingly preferred, with the highest proportion of accepted simulations at 0.9988 and the highest Bayes factor at 823. In this model comparison, only simulations from models A and B were accepted. In cross-validation analysis the five models could be differentiated from each other (Table S5).

We note that the weighted medians inferred under the original and revised demographic models for gene flow into the common ancestors of eastern and western gorillas respectively, are highly correlated, as expected (B a

nd D: rho=0.8531903, p=1.075e-06; C and E rho=0.8870695, p=2.913e-06).

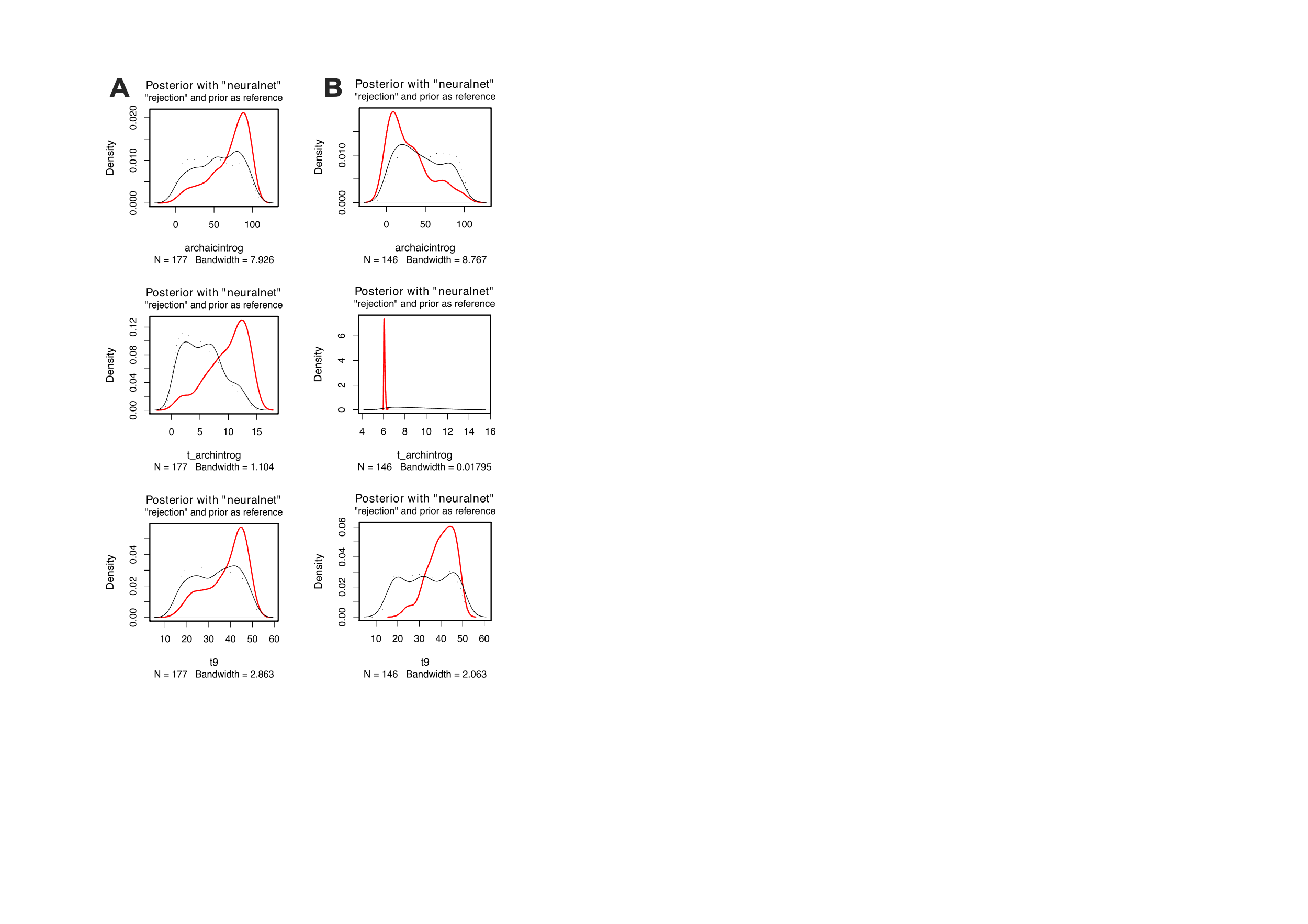

**Figure S11:** Posterior distributions for the archaic introgression proportion, time of archaic introgression, and gorilla-ghost split time, for the revised models of ghost gene flow to **A** the common ancestor of eastern gorillas and **B** the common ancestor of western gorillas, sampling all parameters from priors. We note these are equivalent to Fig 2C, but for the revised ghost models (models D and E). The dotted line indicates the prior distribution. The black line indicates the posterior inferred with a simple ‘rejection’ algorithm. The red line represents the posterior inferred with neural networks. Distributions are plotted in ms units.

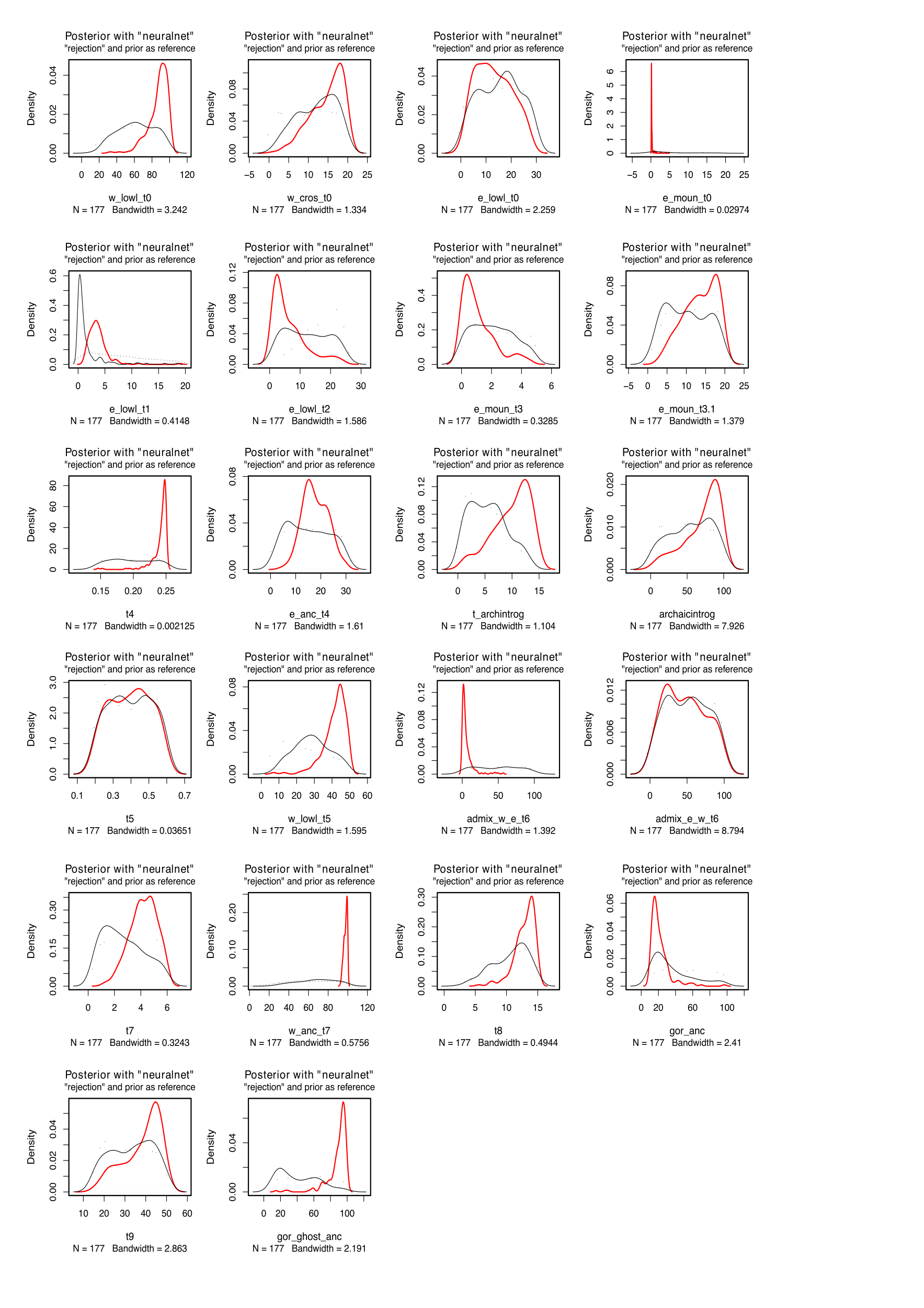

**Figure S12:** Posterior distributions for all parameters inferred under the revised model of ghost gene flow to the common ancestor of eastern gorillas sampling all parameters from priors. Red indicates the posterior distribution inferred with neural networks. Black indicates the posterior distribution inferred under a rejection method. The dotted grey line indicates the prior distribution.

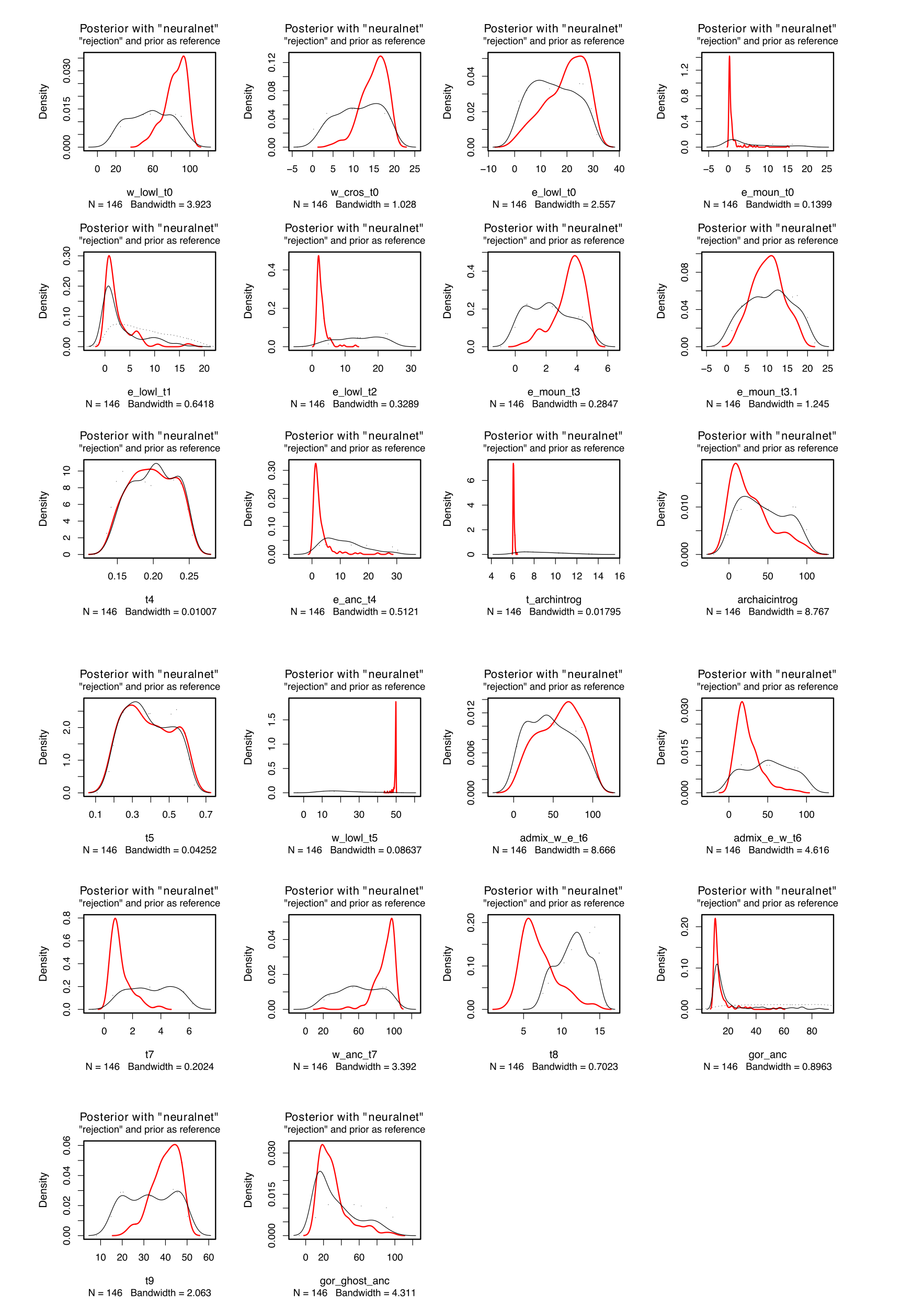

**Figure S13:** Posterior distributions for all parameters inferred under the revised model of ghost gene flow to the common ancestor of western gorillas sampling all parameters from priors. Red indicates the posterior distribution inferred with neural networks. Black indicates the posterior distribution inferred under a rejection method. The dotted grey line indicates the prior distribution.

#### 3.5 Parameters from hmmix

We inferred parameters from the HMM model in hmmix, as shown in Table S11. The coalescence times between the two gorilla species are inferred at ~256 kya, which is more recent than the estimates from the ABC modelling. However, reversing ingroup and outgroup (i.e. using western lowland gorillas as potential ingroup and eastern gorillas as potential outgroup) yields a larger coalescence time of ~572 kya due to the larger effective population size. This relationship between population size and coalescence time makes it difficult to compare to the divergence times of the ABC modelling. Furthermore, we infer a coalescence time of gorilla and ghost segments in eastern gorillas at 1,520 Mya. However, the archaic percentage is inferred at 17.7%, which then represents a larger archaic proportion at a shallower coalescence. The calculated admixture time is ~69 kya, hence older than the one inferred in the demographic model as well. A thorough filtering for decoding the introgressed fragments with hmmix (Methods) leads to a largely overlapping set of candidate regions.

#### 3.6 Validation of method performance

To assess the performance of the *S** statistic and hmmix and their robustness to demographic model misspecifications we performed validation analyses, following the approach of (Huang et al. 2022). This is particularly pertinent for the *S** statistic, which requires a null demographic model (without ghost introgression) to determine outliers of the statistic.

We generated simulations using msprime (Baumdicker et al. 2022; Kelleher, Etheridge, and McVean 2016) under different null demographic models and assessed the performance of the *S** statistic and hmmix using precision-recall curves (Fig S14). We define precision as the number of true introgressed fragments of all introgressed fragments inferred, and recall as the number of inferred true introgressed fragments of all true introgressed fragments (ie recall represents the detection rate of true introgressed fragments) (Huang et al. 2022).

We simulated data and generated general linear models of the expected distribution of *S** scores under 1) the ABC-based null demographic model (Fig S6A) which is the ‘main model’ here and 2) the ‘worst null model’, where we take the maximum value of the 95% credible interval for all ancestral Ne parameters (rather than the weighted median posteriors), this hence increases the number of highly divergent haplotypes present due to incomplete lineage sorting (ILS) (rather than introgression) (Table S3).

We then simulated data under the model of archaic introgression into the eastern ancestor (model B), as well as a modified model of archaic introgression with the maximum values of the 95% credible interval for all ancestral Ne parameters (“worst” model B). We subsequently run *S** and hmmix, with a range of values for the quantile (threshold to define outliers of the statistic) of 0-0.999 for *S** and the posterior probability of 0-0.9999 for hmmix, following (Huang et al. 2022) (Table S8-S9). For each model we simulated 200 Mb with 100 replicates and sampled 1 individual for the target population (eastern lowland or mountain gorillas) and 10 individuals for the outgroup population (western lowland gorillas).

Additionally, for the *S** statistic we explore a ‘worst mis-specified’ scenario, where we generate simulated data under the ‘worst model’ (with high ILS) but run the *S** analysis using the outlier values inferred for model A (expecting less ILS) (Fig. S14).

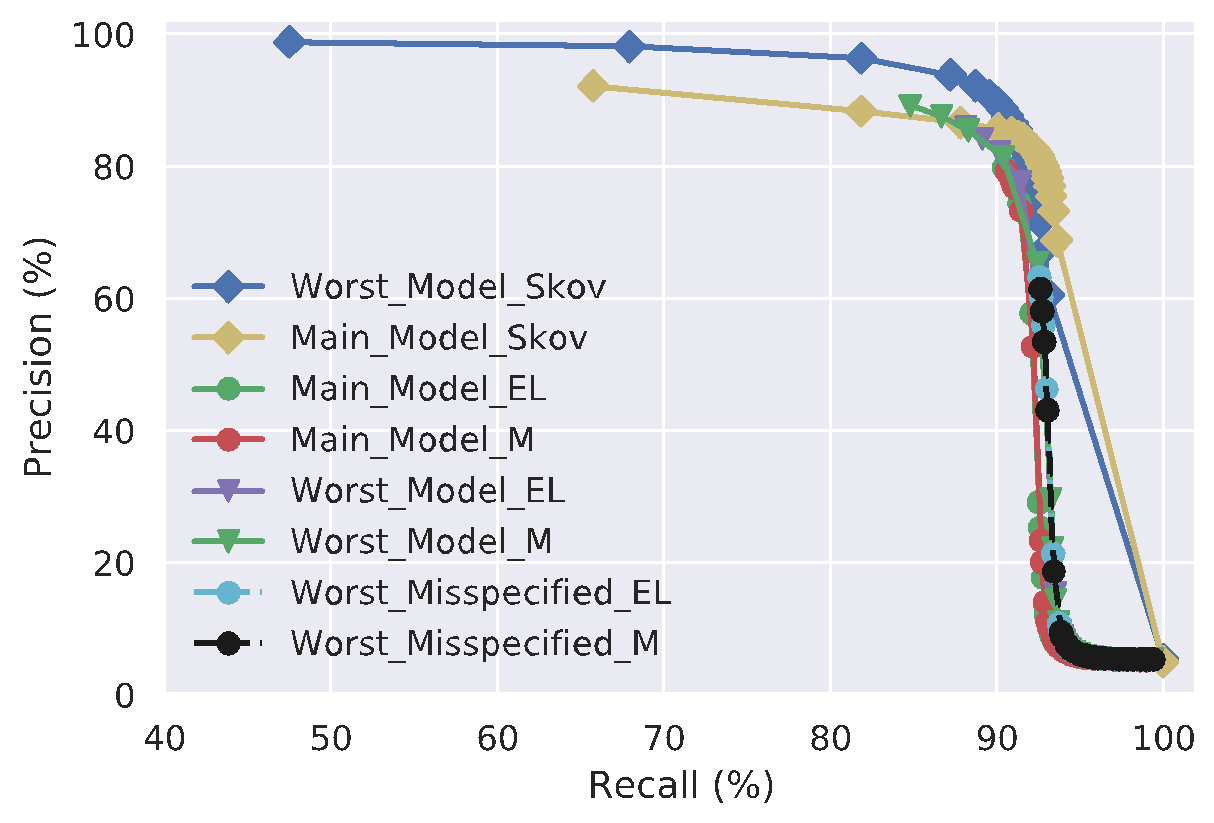

**Figure S14:** Precision-recall curves under different null demographic scenarios for the *S** statistic at the 99% quantile as implemented in sstar (Huang et al. 2022) and hmmix at the 95% posterior probability cutoff. Main model refers to a model taking the weighted median posteriors from the ABC-based null demography presented herein (Fig S6A). Worst model refers to a model taking the maximum value of the 95% credible interval for all ancestral Ne parameters from the ABC-based null demography. For the *S** statistic we consider the target population as alternately eastern lowland or mountain gorillas, eg Main Model EL. Worst mis-specified is where we generate simulated data under the worst model but run the *S** analysis using the ‘quantile’ or outlier values inferred under the main model. Skov=hmmix method, EL=eastern lowland gorillas, M=mountain gorillas.

### 4. Characterising introgressed fragments

#### 4.1 General features

*S** scores are correlated with the numbers of segregating sites, as expected (Fig. S15); putatively introgressed windows are observed for high *S** scores across this distribution, depending on the demographic model.

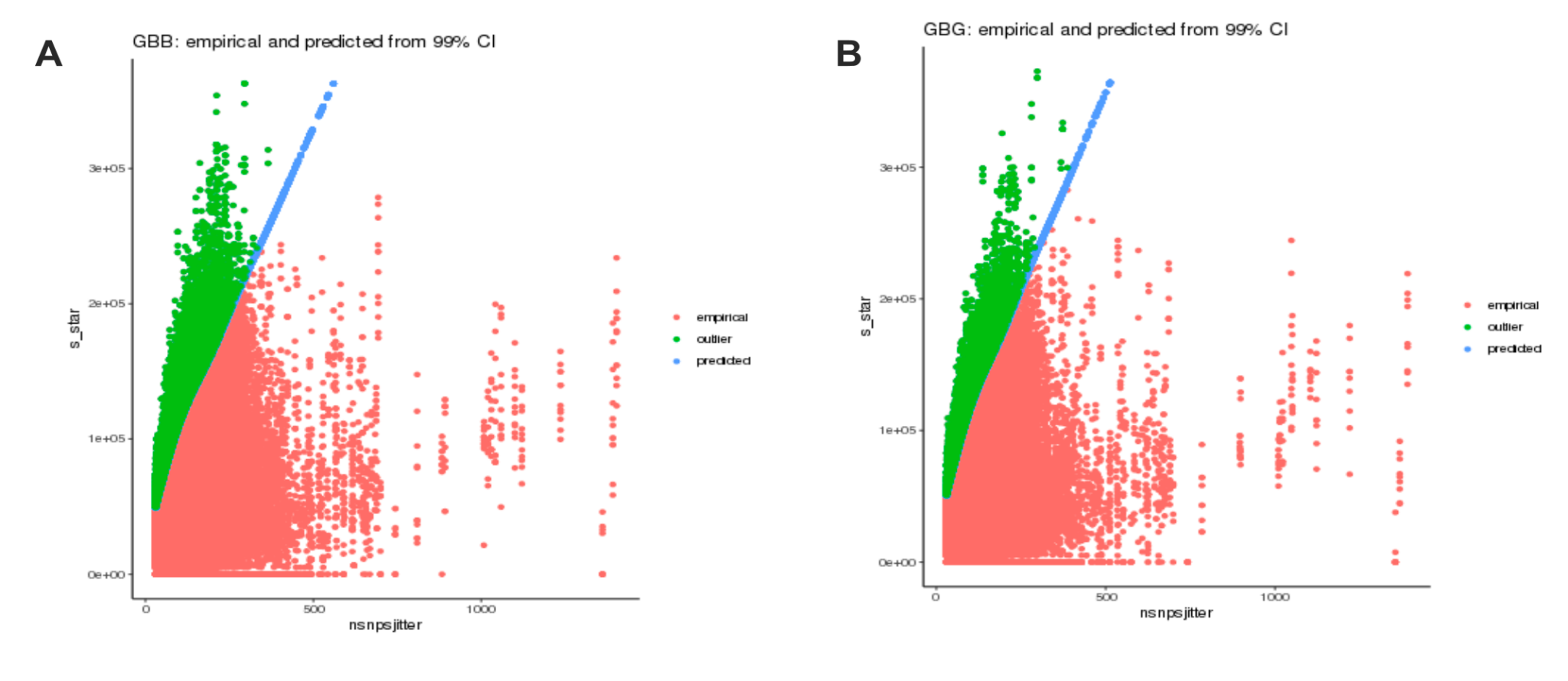

**Figure S15:** Genome-wide distributions of *S** scores calculated in 40kb windows for **A** mountain gorillas as ingroup, western lowland gorillas as outgroup, **B** eastern lowland gorillas as ingroup, western lowland gorillas as outgroup. Red indicates the genome-wide distribution, blue the predicted *S** values under the general linear model (Methods) and green the outlier windows inferred under the 99% confidence interval.

We do not observe a significant difference in introgressed fragment length distributions between the eastern subspecies, as inferred under hmmix (p>0.01, Wilcoxon unpaired test for both 0.9 and 0.95 threshold) (Fig S16).

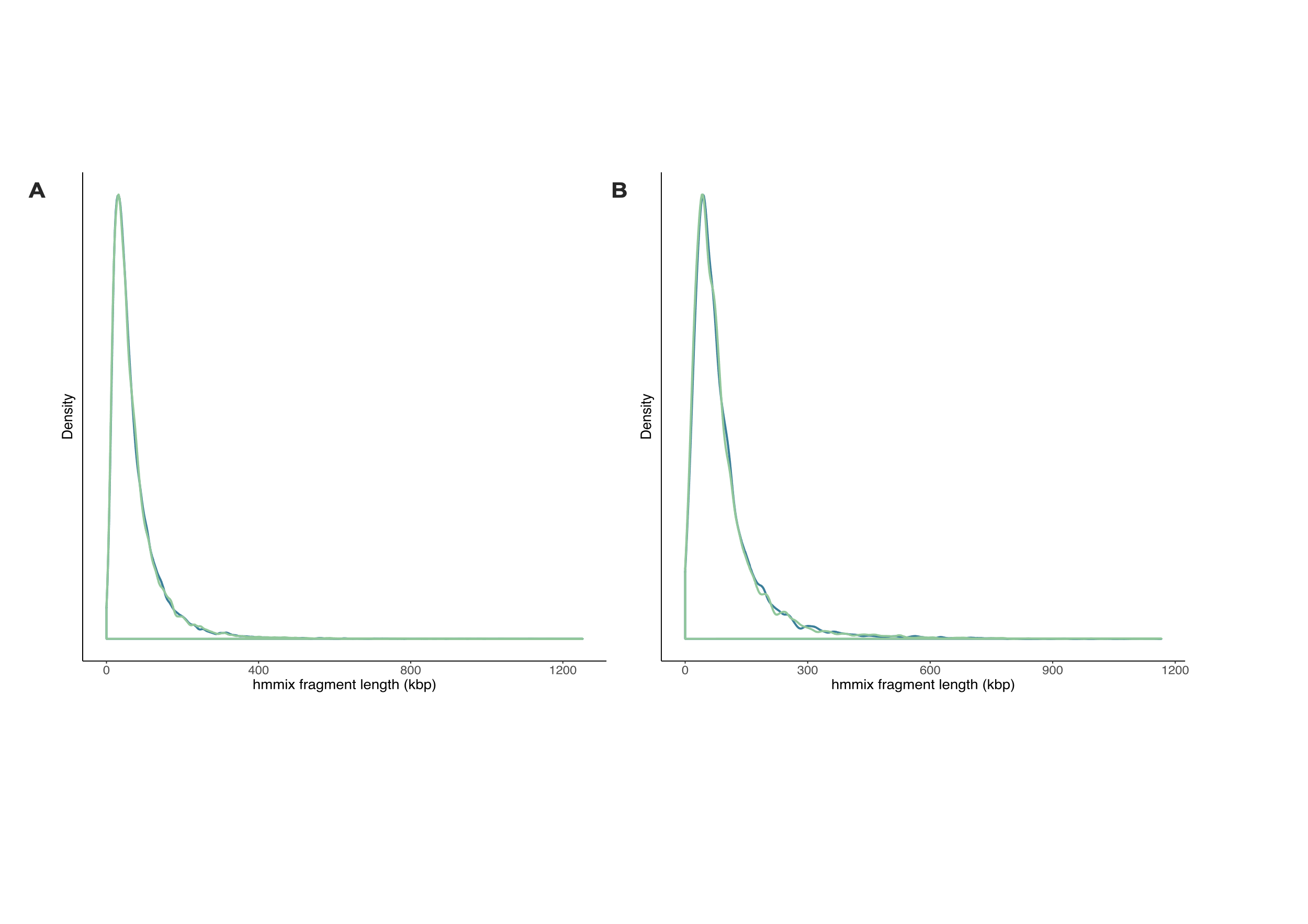

**Figure S16:** Distribution of hmmix fragment lengths for mountain gorillas (blue) and eastern lowland gorillas (green) at a threshold of A 0.9 and B 0.95.

When we compare a PCA of putative introgressed regions against a PCA of random regions of equivalent length distribution, we see in the introgressed regions that PCs 1 and 2 explain a greater percentage of the variance (Fig. S17). For introgressed regions PC1 exhibits increased separation of eastern from western gorillas, as expected under archaic introgression specifically into eastern gorillas (Kuhlwilm et al., 2019). While PC2 separates out the eastern subspecies to a greater extent than in random regions. We note that in PC1 the target individual carrying the introgressed material tends to fall outside the variation of its subspecies, but this is not always the case, indicative of the population frequency of the introgressed regions. Likewise in phylogenetic trees the target individual carrying the introgressed material has a longer branch, rather than falling basal to the other sequences (Fig S18). Haplotype networks of putatively introgressed regions often show expected patterns (Fig. S19), where the putatively introgressed haplotype shows an unusually large divergence to the variation observed among gorillas. Among the 20 longest introgressed regions, 90% of the resulting haplotype networks look archaic in origin.

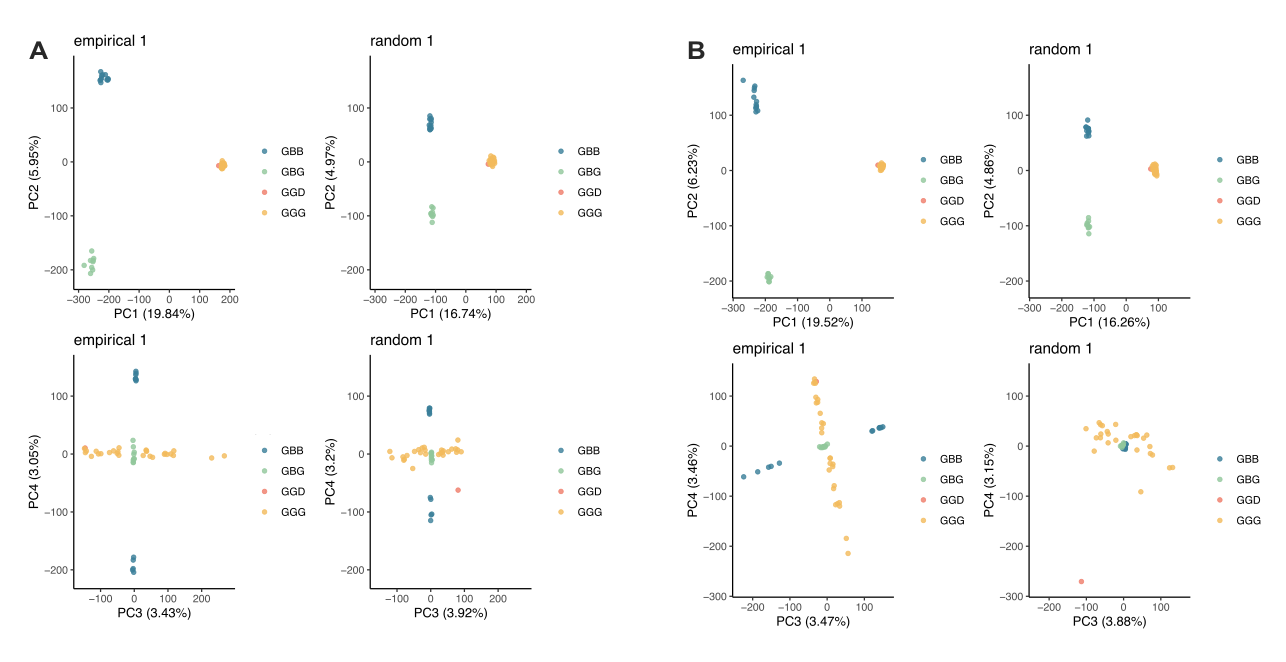

**Figure S17: A** PCA of SNPs in the putative introgressed regions of eastern lowland individual 1 (*Gorilla_beringei_graueri-9732_Mkubwa)* and of random genomic regions of equivalent length distribution. PCs 1-4 are shown. **B** Equivalent PCA analysis for putative introgressed regions of mountain gorilla individual 1 (*Gorilla_beringei_beringei-Bwiruka).*

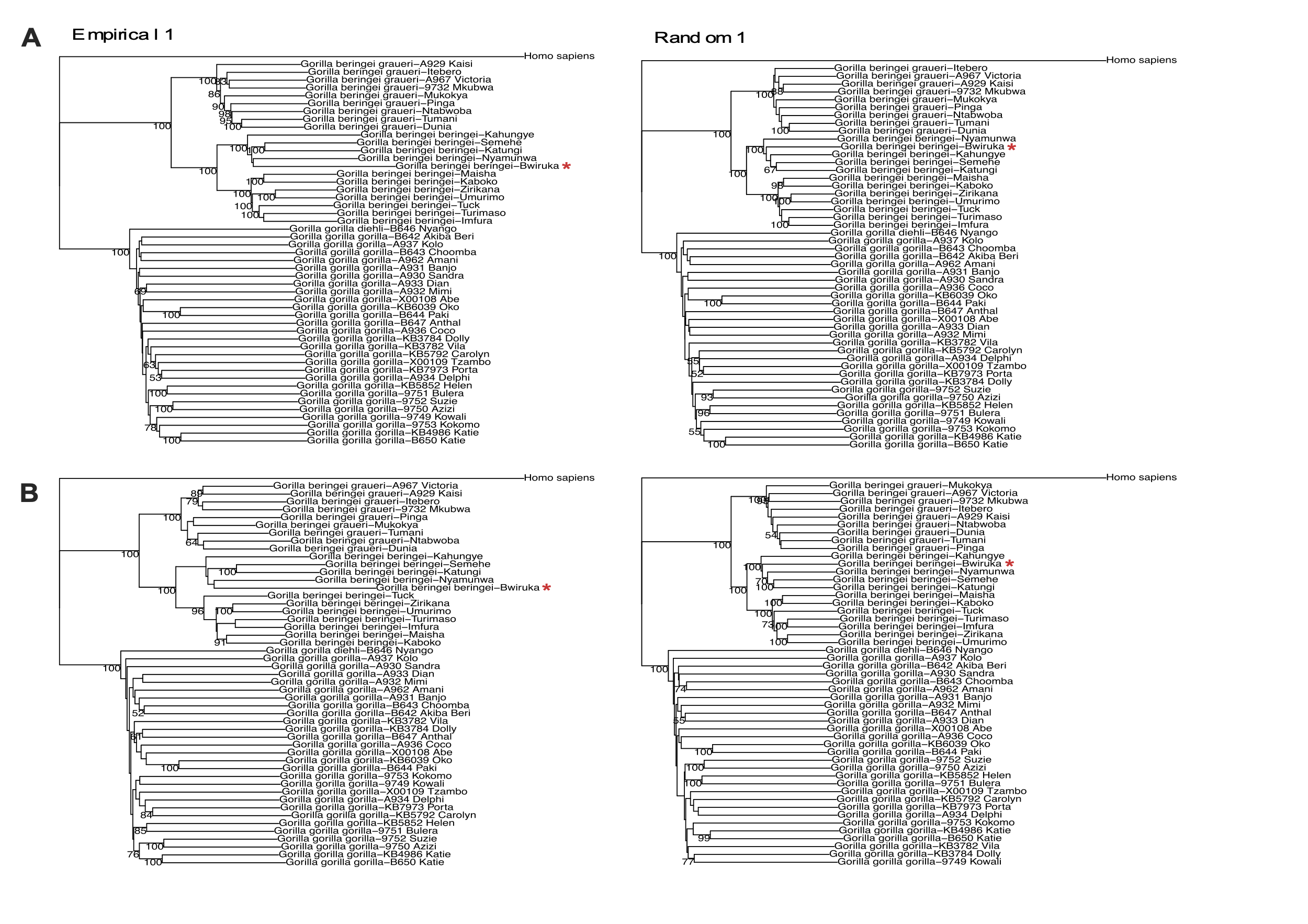

**Figure S18: A** NJ tree of SNPs in all putative introgressed regions of mountain gorilla individual 1 (*Gorilla_beringei_beringei-Bwiruka)* and of random genomic regions of equivalent length distribution. **B** NJ tree of SNPs in putative introgressed regions unique to mountain gorilla individual 1 (so-called ‘private introgressed regions’) and equivalent random genomic regions. The target individual is indicated by a red star.

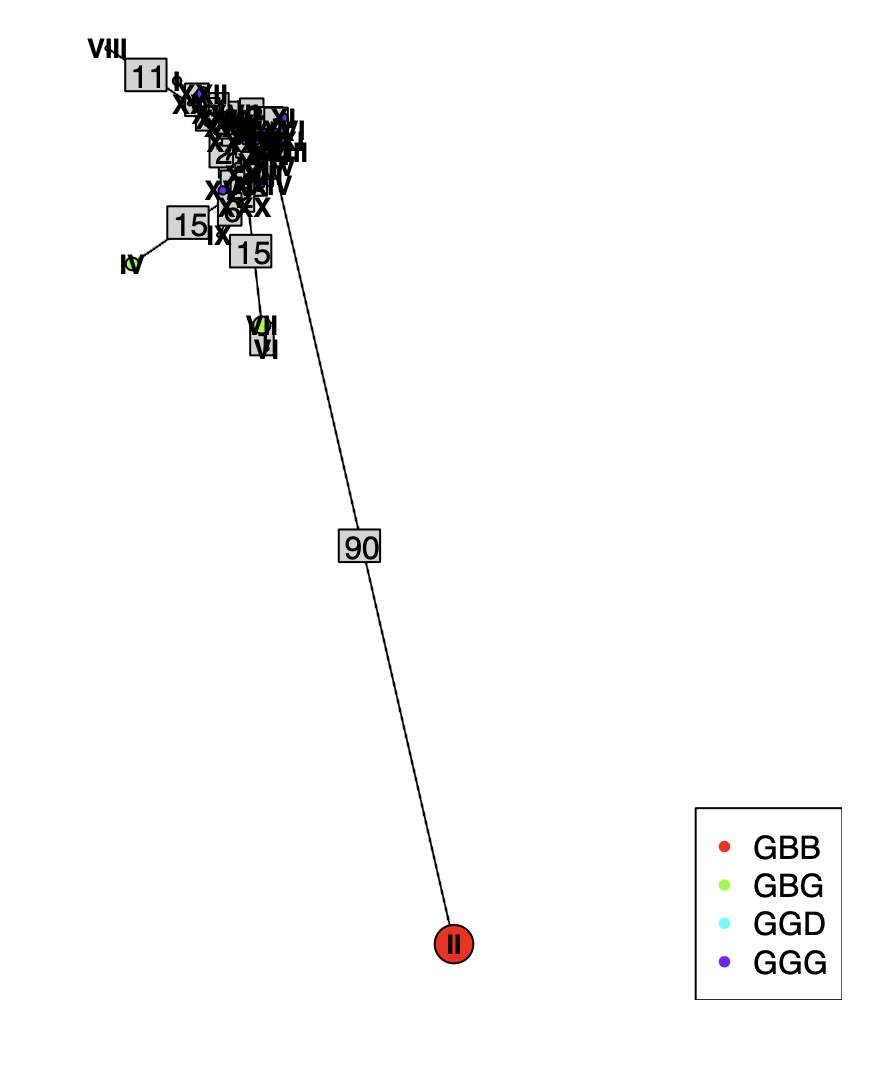

**Figure S19:** Haplotype network of one of the putative introgressed regions (chr13: 79839000-80119000), which looks characteristic of archaic introgression, where haplotype II carried by mountain gorillas is far outside the diversity of other gorillas.

#### 4.2 Introgression and selection

We find no depletion in the proportion of protein-coding base pairs (bp) in the introgressed regions compared to random regions of the genome (Fig. S20). Putative deserts of introgression of more than 5 Mbp are rare (Fig. S21). In Fig. S22, we show the likelihood scores for candidate genes for adaptive introgression, using VolcanoFinder.

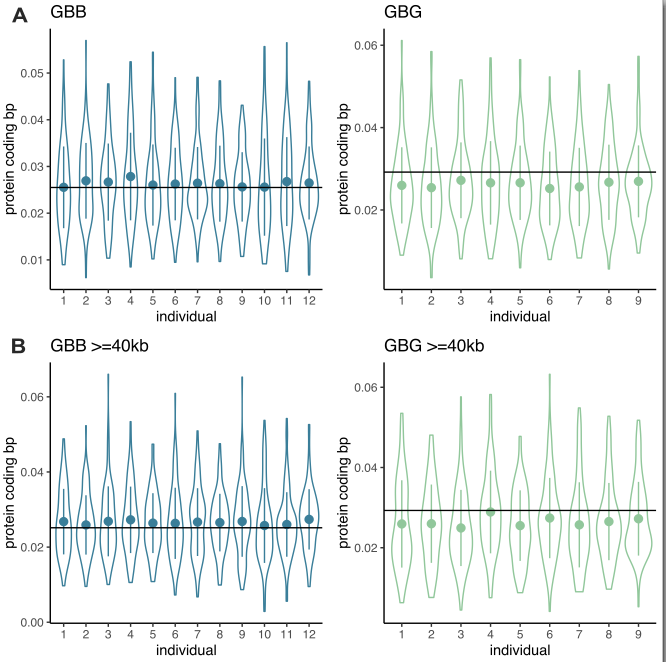

**Figure S20: A** Proportion of protein coding base pairs in putative introgressed regions (lines) and in random genomic regions (violin plots) per individual for both eastern gorilla populations. **B** Proportion of protein coding base pairs in putative introgressed regions of length >= 40kb (lines) and in equivalent random genomic regions (violin plots) per individual.

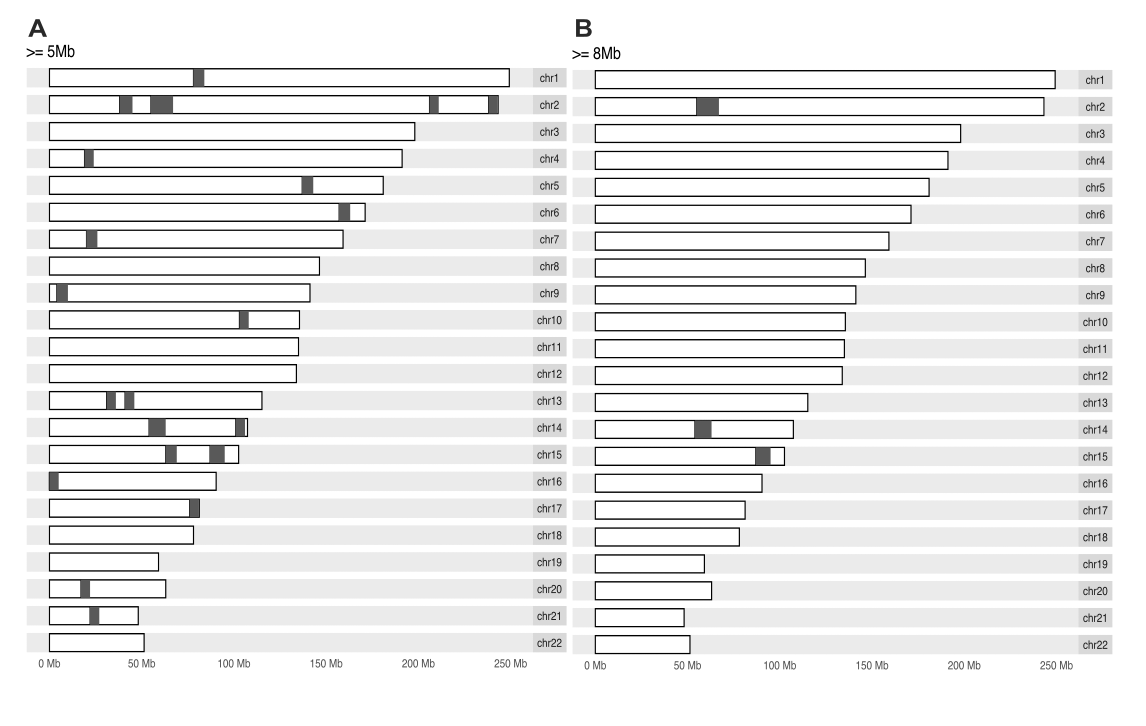

**Figure S21:** Distribution of regions depleted of archaic introgression **A** regions of length >=5Mb **B** regions of length >=8Mb.

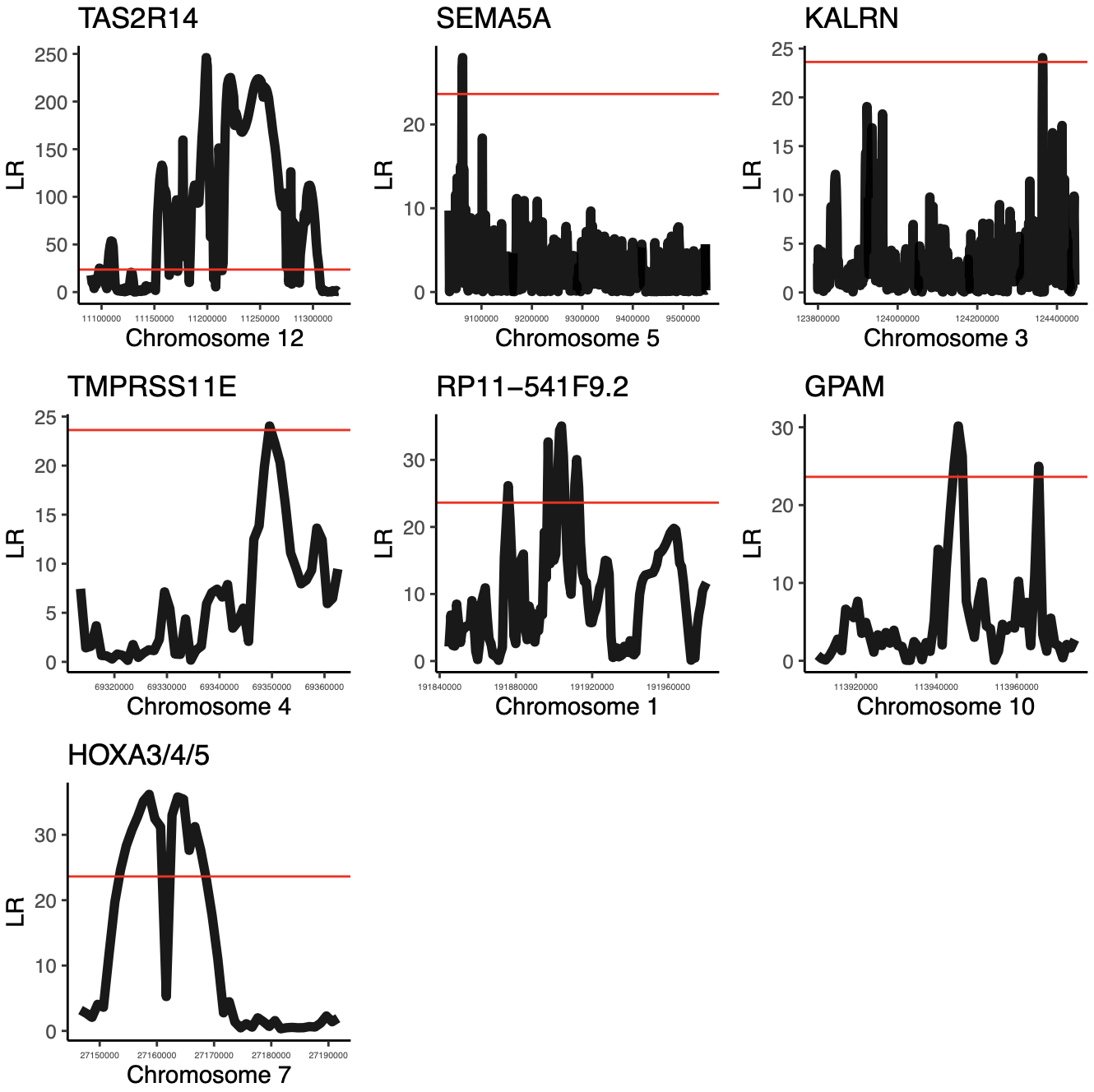

**Figure S22:** Likelihood ratio scores for the seven candidate genes of adaptive introgression, estimated with VolcanoFinder. Red line indicates the 95% threshold for the VolcanoFinder likelihood ratio score.

#### 4.3 Functional consequences: mutational tolerance

To address the question of mutational tolerance, specifically whether more deleterious mutations are observed in introgressed rather than random genomic regions, we assessed different measures of deleteriousness, using: genomic evolutionary rate profiling (GERP), sorting intolerant from tolerant (SIFT), polymorphism phenotyping (PolyPhen-2) and LINSIGHT scores (Davydov et al. 2010; Kumar, Henikoff, and Ng 2009; Adzhubei et al. 2010; Huang, Gulko, and Siepel 2017). We downloaded the pre-computed base-wise GERP scores for hg19 (Davydov et al. 2010) and considered sites (>4) as having high functional impact and sites (-2<x<2) as having low or likely neutral impact. SIFT and PolyPhen-2 scores were extracted from VEP annotation for missense variants. We consider sites annotated with (SIFT=‘deleterious’ or ‘deleterious_low_confidence’; PolyPhen-2=‘probably_damaging’ or ‘possibly_damaging’) as high impact and (SIFT=‘tolerated’ or ‘tolerated_low_confidence’; PolyPhen-2=’benign’) as low impact. LINSIGHT scores incorporate epigenomic information, including chromatin accessibility and transcription factor binding (Huang, Gulko, and Siepel 2017). We downloaded the pre-calculated LINSIGHT scores for hg19 (Huang, Gulko, and Siepel 2017).

For GERP, SIFT and PolyPhen-2 scores we calculated the proportion of high impact sites within putative introgressed regions and random regions of equal length distribution and sufficient callable sites (high / high and low impact sites). We calculated the mean LINSIGHT score across regions, since few high impact sites (>0.8) were identified in our dataset. We find a higher proportion of high impact GERP sites in introgressed regions of eastern lowland gorillas compared to mountain gorillas (Fig. 3E). However, for SIFT, PolyPhen-2 and LINSIGHT scores the introgressed regions of both eastern lowland and mountain gorillas follow random expectation (Fig. S23).

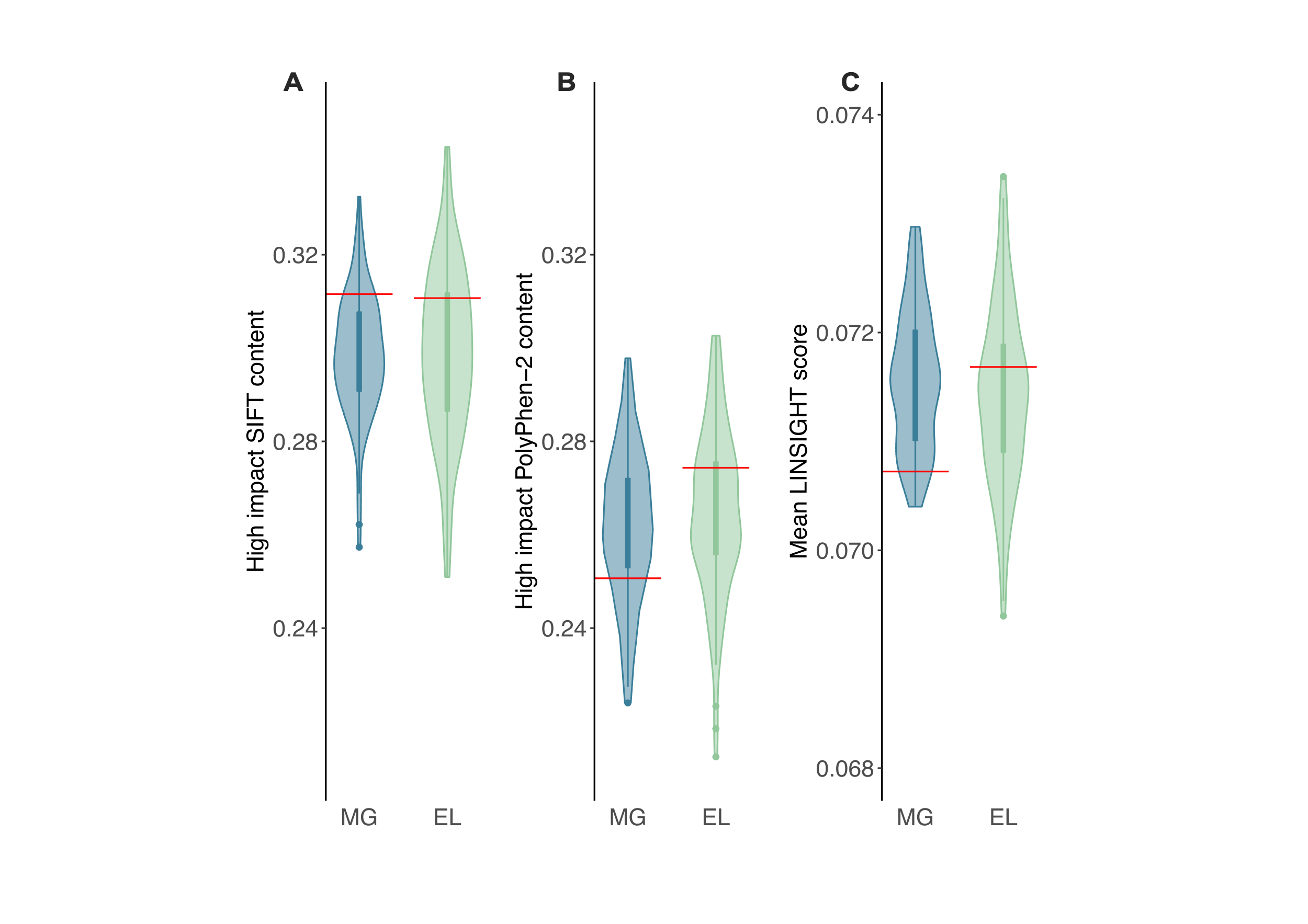

**Figure S23:** Mutational conservation in introgressed fragments. Proportion of high impact sites in introgressed regions (red lines) and random regions (violin plots) for **A** SIFT scores and **B** PolyPhen-2 scores. High impact sites are those annotated as ‘deleterious’ and ‘deleterious low confidence’ for SIFT, and ‘probably damaging’ and ‘possibly damaging’ for PolyPhen-2. **C** Mean LINSIGHT score across introgressed regions (red lines) and random regions (violin plots). In panels A-C MG = mountain gorillas, EL = eastern lowlands.

#### 4.4 Functional consequences: regulatory elements

We undertook an investigation of regulatory elements in introgressed fragments. We assessed the proportion of regulatory base pairs within putative introgressed and random regions of equivalent length and callability, using the gorilla-defined regulatory element annotations of García-Pérez et al. (García-Pérez et al. 2021). We assess this both from a global perspective and per regulatory element type (poised, strong, weak, enhancers and promoters). To do this, we performed a sequential liftover of the coordinates from gorGor4 to hg38 to hg19, to match the genomic coordinates used for the rest of our analyses. We filtered out entries corresponding to non-regulatory elements in at least one of the two replicates of gorilla lymphoblastoid cells (Non-re, E/Non-re and P/Non-re). We also filtered out entries annotated as ambiguous (aE, aP, P/E).

We note that this data derives from gorilla lymphoblastoid cells (LCLs), which means that the patterns of expression may be cell-type dependent and specific regulatory effects, for example during brain development, would not be recovered. This is an inherent limitation of this kind of analysis in a non-human context. Moreover, the two gorilla LCL replicates belonged to the western species, hence are equidistant to both eastern subspecies.

We assessed the gene regulatory architecture of strong enhancers (sE) in mountain gorilla introgressed regions and equivalent random genomic regions, using the definitions of García-Pérez et al. 2021. We considered enhancer-interacting enhancers, intragenic enhancers, enhancers within promoters (genic promoters), promoter-interacting enhancers and proximal enhancers (EiE, gE, gP, PiE, prE). This analysis is restricted to sE associated with genes.

We find no difference in the overall proportion of regulatory base pairs in putative introgressed regions compared to random genomic regions for either eastern gorilla population (Fig. S24A). However, when we consider the proportion of regulatory base pairs per regulatory element we see an excess of sE in mountain gorilla introgressed regions, compared to random regions (Fig. S24B). These sE are largely intragenic enhancers (Fig. S25), which agrees with patterns of regulatory architecture observed in primate sE more generally by (García-Pérez et al. 2021).

Furthermore, García-Pérez et al. (García-Pérez et al. 2021) had annotated which genes are associated with each regulatory element. Taking these annotations and filtering to genes with one-to-one orthologs across the primates considered by García-Pérez et al. (García-Pérez et al. 2021) (humans, chimpanzees, gorillas, orangutans and macaques) we define two sets of candidate genes: 1) genes regulated by sE in mountain gorilla introgressed regions (235 genes), and 2) genes regulated by sE in mountain gorilla adaptively introgressed regions (45 genes) (Tables S18-S19). We performed an over-representation analysis of our candidate genes for gene ontology terms using the WebGestaltR package and default settings (Liao et al. 2019). Our background set consisted of genes regulated by gorilla sE (again taking those genes with one-to-one orthologs in primates) (Table S20). No gene ontology category reached the significance threshold of FDR=0.05 with Benjamini-Hochberg correction (Fig. S26). The top gene ontology categories detected relate to the LCL cell type, namely ‘establishment of lymphocyte polarity’ (p-value=0.00018256, FDR=0.38985) and ‘establishment of T cell polarity’ (p-value=0.00018256, FDR=0.38985) for candidate gene set 1) and ‘forebrain generation of neurons’ (p-value=0.000066791, FDR=0.14263) for candidate gene set 2).

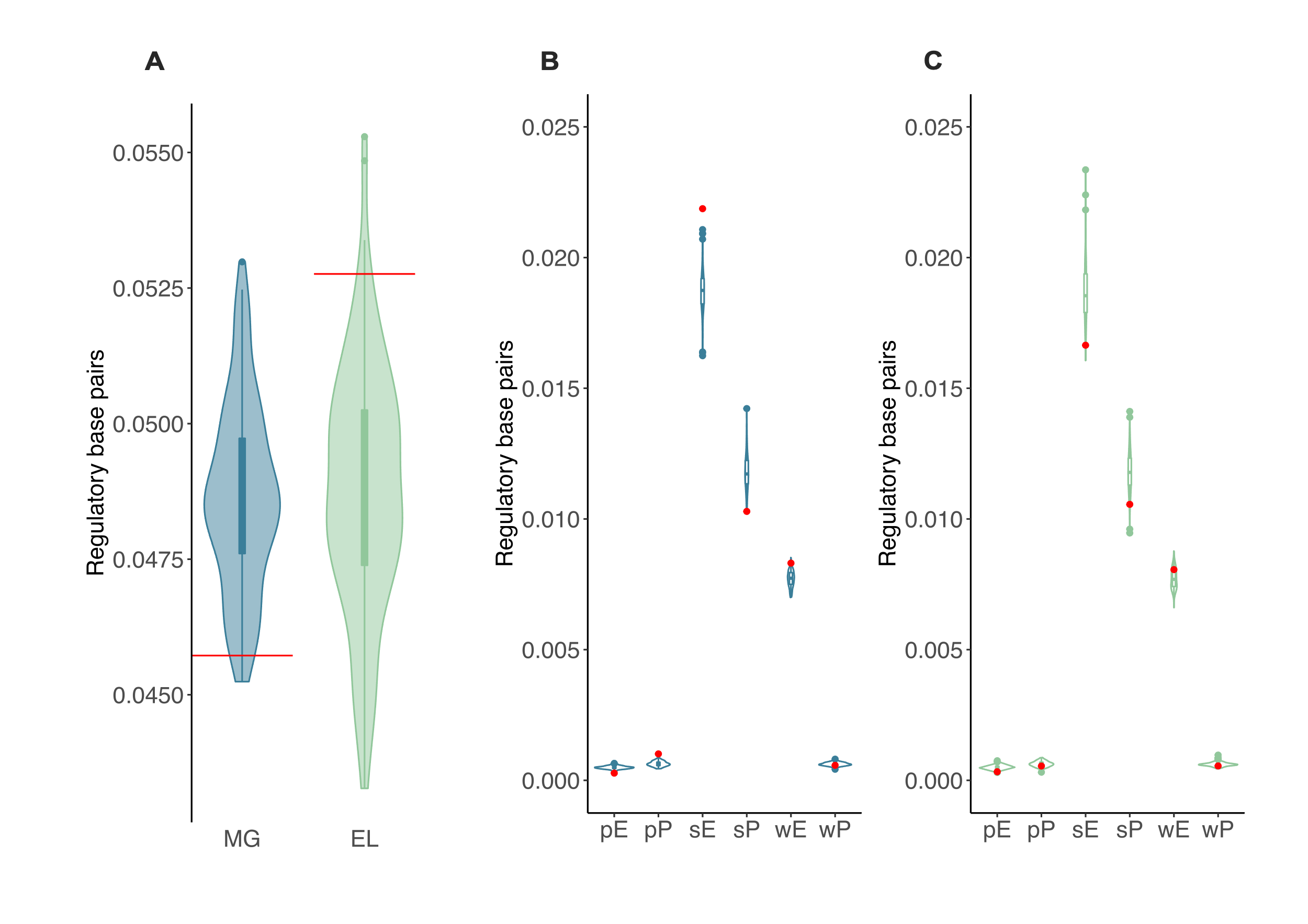

**Figure S24:** Proportion of regulatory base pairs in introgressed regions (red lines) and random regions (violin plots) population wide in **A** and per regulatory element type for **B** mountain gorillas and **C** eastern lowland gorillas. Abbreviations represent: pE=poised enhancer, pP=poised promoter, sE=strong enhancer, sP=strong promoter, wE=weak enhancer, wP=weak promoter. In panel A MG = mountain gorillas, EL = eastern lowlands.

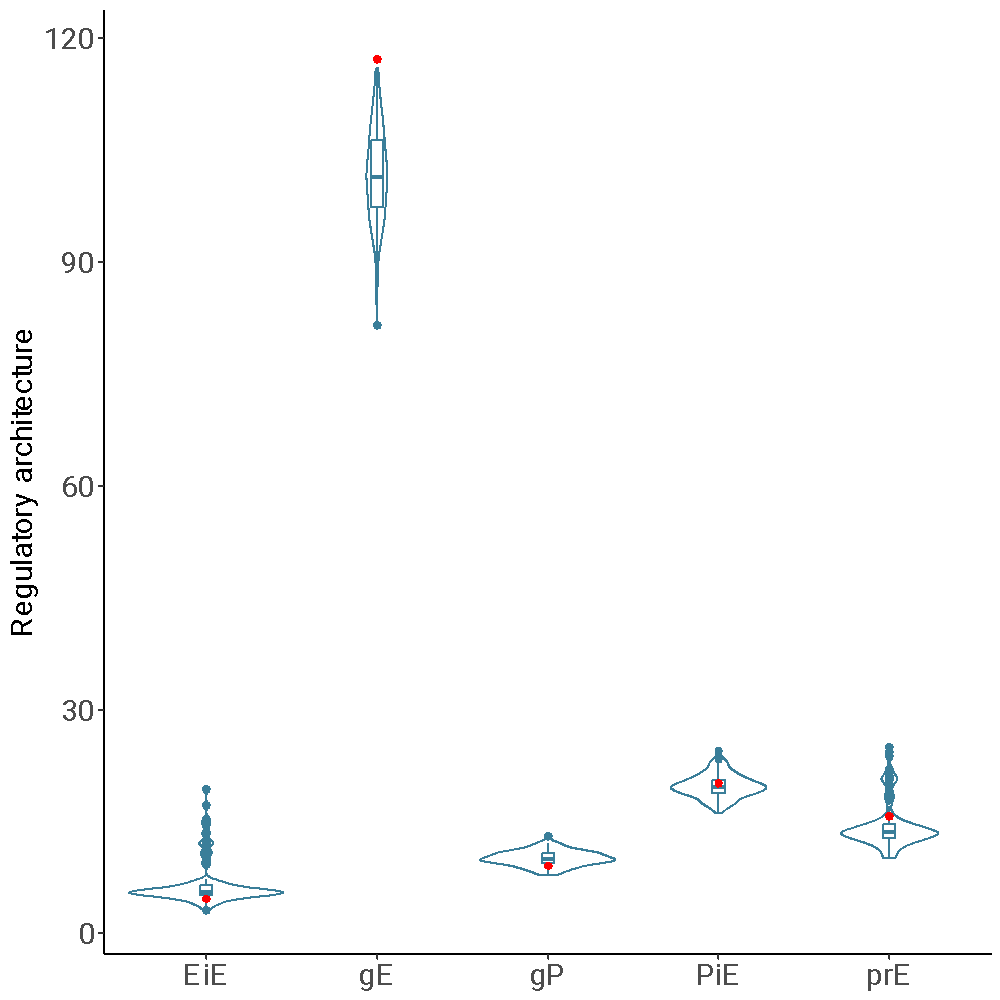

**Figure S25:** Gene regulatory architecture of strong enhancers in mountain gorilla introgressed regions (red points) and random genomic regions of equivalent length and callability (violin plots). Abbreviations represent: EiE=enhancer-interacting enhancer, gE=intragenic enhancer, gP=genic promoter, PiE=promoter-interacting enhancer, prE=proximal enhancer. This analysis only considers those strong enhancers which could be annotated to genes by (García-Pérez et al. 2021).

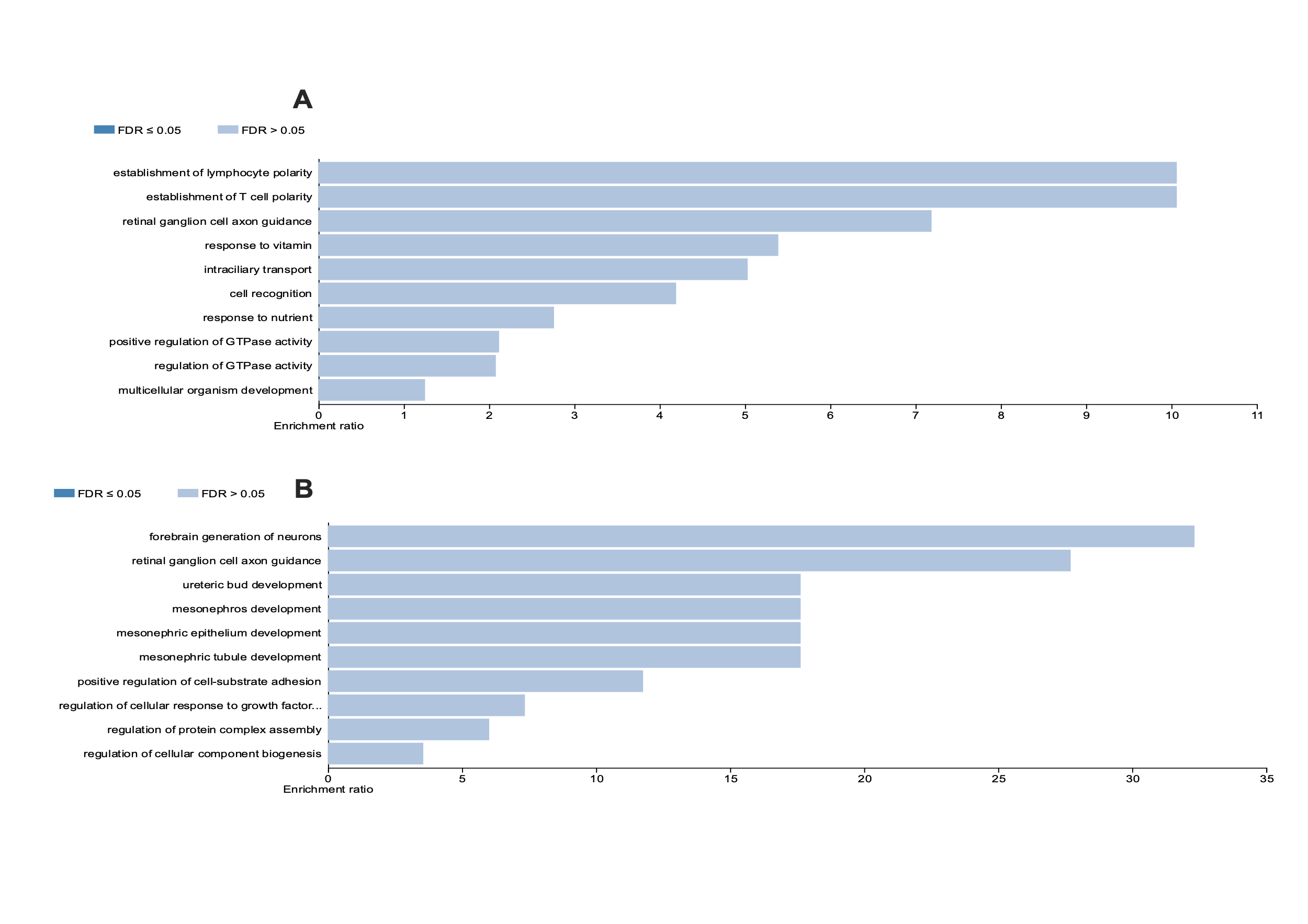

**Figure S26:** Over-representation in gene ontology categories for **A** genes regulated by sE in mountain gorilla introgressed regions and **B** genes regulated by sE in mountain gorilla adaptively introgressed regions. No category reaches significance at FDR=0.05 with Benjamini-Hochberg correction.
